# Sex differences in neural circuits driving binge drinking: A female-specific role for an amygdalo-striatal pathway

**DOI:** 10.64898/2026.02.10.705193

**Authors:** Xavier J Maddern, Amy J Pearl, Qian T Tan, Harry Dempsey, Lauren T Ursich, Kade L Huckstep, Brandon K Richards, Roberta G Anversa, Erin J Campbell, Andrew J Lawrence, Robyn M Brown, Leigh C Walker

## Abstract

**Background:** Rates of binge drinking have converged significantly between the sexes over recent decades, driven by increased rates of alcohol misuse in women. However, understanding of fundamental circuitry and neurobiology driving alcohol use in females, or how this may differ from male subjects remains underexplored.

**Methods:** We quantified c-Fos expression across 40 brain regions in alcohol naïve, alcohol anticipating and binge drinking male and female mice. *In vivo* fiber photometry examined sex differences in basolateral amygdala (BLA) activity changes to alcohol intake. Chemogenetic BLA inhibition investigated a functional role in binge drinking. We then assessed sex differences in BLA efferent projection activation following binge drinking. Finally, we functionally interrogated the BLA to nucleus accumbens core (AcbC) projection in binge drinking.

**Results:** Binge drinking reduced network modularity (number of communities with similar activation patterns) in both sexes relative to alcohol naive and anticipating same-sex counterparts. Female binge drinking mice had increased BLA c-Fos expression compared to female naïve and male binge drinking counterparts. *In vivo* fiber photometry revealed greater and more prolonged BLA responsivity at the onset of alcohol intake in females. Global BLA inhibition reduced reward intake in both sexes. However, the BLA to AcbC projection was preferentially activated in female binge drinking mice, and inhibition of this pathway reduced binge alcohol intake exclusively in females.

**Conclusions:** We identified sex differences in the neural circuits engaged in binge drinking, highlighting the BLA to AcbC projection may in part underpin sex differences in alcohol misuse. This provides further evidence of distinct neurobiological drivers of alcohol-related behaviors between the sexes.

## Introduction

Alcohol misuse is a leading risk factor for disability and premature death among individuals aged 20-39 years (1), accounting for ∼5.1% of the global disease burden (2). The most common form of alcohol misuse in adolescents and young adults is binge drinking (3), a robust predictor of the future development of alcohol use disorder (AUD) (4, 5). While rates of alcohol use, misuse and AUD have historically been higher in men than in women, these rates have converged significantly in recent decades, driven by increased alcohol use in women (6-8). Of concern, women are more vulnerable to alcohol-induced health consequences (9), transition faster to problematic drinking and AUD (10, 11), and are more likely to consume alcohol to cope with negative affect (12). Despite this, recognition of sex as a biological variable has accelerated only in recent years, leaving critical gaps in our understanding of female biology (13, 14).

The limited clinical literature to date suggests sex differences in neural activation in response to alcohol-associated cues in social and problematic drinkers (15-17), but not in individuals with AUD (18). While there are inconsistent differences reported following exposure to alcohol itself in social drinkers (19, 20). Preclinically, several studies have characterized brain activation in male rodents following high levels of alcohol exposure, with these findings being largely in agreement across forced and voluntary administration models (21-27). However, similarities and differences in brain activation between sexes in the anticipation of, and response to, excessive alcohol intake have not been well-established. Recent studies including both male and female rodents have reported network level neurobiological sex differences in mice with a frontloading drinking phenotype (28) and following a 4hr drinking-in-the-dark (DID) session (29). Others have found discrete sex differences in the involvement of select neuropeptides, receptors and cell populations within (30-35), and pathways from (36), specific loci in excessive alcohol intake.

Notably, no studies to date have examined activity patterns throughout the brain driven by anticipation of alcohol, and compared to excessive alcohol consumption itself, to assess neurobiological sex differences at both a network and circuit level. Here, we found reduced network modularity (number of communities with similar activation patterns) following binge drinking in both sexes relative to alcohol naïve and anticipating same-sex counterparts. We observed a sex-specific increase in basolateral amygdala (BLA) c-Fos expression in female binge drinking mice, confirmed by *in vivo* fiber photometry. While chemogenetic BLA inhibition led to generalized reductions in reward intake, we identified preferential activation of the BLA to nucleus accumbens core (AcbC) projection in female compared to male binge drinking mice. We then found inhibition of the BLA→AcbC projection consistently, but modestly, reduced binge alcohol intake sex-specifically in female mice. Together, these data provide important insights into neurobiological sex differences in different alcohol-related states, and identifies the BLA→AcbC projection as having a sex-specific role in binge drinking in female mice.

## Methods and Materials

For full details please refer to the Supplemental Materials.

### Assessing sex differences in brain activity of alcohol naïve, alcohol anticipating and binge drinking mice

C57BL6/J mice were divided into three groups; alcohol naïve, alcohol anticipating and binge drinking. The naïve group (*n*=7/sex) experienced no experimental manipulation. The alcohol anticipating (*n*=6/sex) and binge drinking (*n*=6/sex) groups completed 10 x 2hr sessions of a binge drinking paradigm (10% v/v ethanol; Fig. S1A) (30, 31). Mice then underwent a final binge drinking session; mice in the alcohol anticipating group received a water bottle in place of the expected ethanol bottle, whilst the binge drinking group undertook a standard drinking session. All mice were perfused immediately following the session (or equivalent time) and brains processed for c-Fos immunohistochemistry (37).

*In vivo* fiber photometry was conducted to further probe BLA responsivity to alcohol. Mice (*n*=6/sex) underwent binge drinking training (6 x 2hr sessions) prior to stereotaxic surgery, where the calcium indicator GCaMP6m was unilaterally microinjected into the BLA and a fiber optic cannula implanted above. Following recovery, mice underwent 8 x 15min recording sessions with ethanol, followed by 2 x 15min sessions to assess natural reward (5% w/v sucrose) induced activity. For all experiments, site validations were confirmed via histology and mice with misplaced injections or cannula were excluded from analysis.

### Examining functional involvement of the BLA in binge drinking

Male (*n*=17) and female (*n*=18) mice underwent stereotaxic surgery to bilaterally microinject an inhibitory Designer Receptor Exclusively Activated by Designer Drugs (DREADD; *n*=10/sex) or control virus (*n*=7 males/8 females) into the BLA during binge drinking training. Following recovery and re-training, mice were tested in an extended 4hr binge session under each treatment condition (vehicle (saline) and clozapine-N-oxide (CNO), 3mg/kg, i.p.), in a randomized counterbalanced manner, with stable baselines reached between tests. Mice were then trained to consume 5% w/v sucrose and tested under identical experimental parameters to assess natural reward consumption.

### Assessing sex differences in activation of BLA outputs during binge drinking

Mice (*n*=24/sex) were trained to binge drink and then underwent stereotaxic surgery to unilaterally microinject cholera toxin subunit β conjugate(s) (CTβ) into one, or two, of the following brain regions; medial prefrontal cortex (mPFC), AcbC, nucleus accumbens shell (AcbSh), bed nucleus of the stria terminalis (BNST) and/or ventral hippocampus (vHipp). Following recovery, mice completed three binge drinking sessions and were transcardially perfused immediately after the last session. CTβ and c-Fos expression were quantified to examine sex differences in activation of BLA outputs following binge drinking (38).

### Examining a functional, sex-specific role for BLA→AcbC pathway in binge drinking

Male (*n*=7) and female (*n*=9) mice were bilaterally microinjected with a retrograde Cre-carrying virus into the AcbC and a Cre-dependent inhibitory DREADD into the BLA. Following recovery and training mice were tested for alcohol and sucrose intake following CNO administration to determine the role of BLA→AcbC projections in binge drinking.

### Statistical analysis

Data were analyzed by *student* t-test, two- or three-way ANOVA’s, or Pearson’s correlations using GraphPad Prism 10. *Post-hoc* fiber photometry analysis was performed using the FiPhoPHA analysis package (39) and brain network analyses using the Python implementation of the Brain Connectivity Toolbox (https://github.com/H-Dempsey/Brain_Network_Analysis_X_Maddern_2026).

## Results

### Sex differences in brain network and regional activation across alcohol-related states

Given the growing evidence of sex differences in the neurobiology of alcohol-related behaviours (40), we explored how alcohol anticipation and binge drinking alter neuronal activation at both the individual locus and network level between sexes (Fig. S1A). Alcohol intake was greater in females compared to males during binge drinking training (main effect of sex, *p*<0.001; Fig. S1B), while no difference was observed in the final 2hr session (*p*>0.05; Fig. S1C; see Table S2 for complete statistical analysis).

We first considered the broader network level changes that may be influencing behavior (41), constructing undirected, weighted and signed inter-regional correlation matrices for each treatment group (Fig. S2). Louvain community detection and consensus clustering was then performed to detect communities/modules of regions with similar patterns of activation (Fig. 1). Assessment of network modularity, number of communities/modules with similar patterns of activation (42), revealed male alcohol naive and alcohol anticipating groups had 4 modules, while the male binge drinking group had 2 modules. Similarly, the female alcohol naive and alcohol anticipating groups had 3 modules compared to the female binge drinking group which had 2 modules. For each treatment group, we calculated the mean coactivation, anatomical coactivation (globally and of anatomical divisions), and community coactivation (Tables S3-4). To statistically compare these metrics between groups, we quantified the empirical difference between the two groups and generated a null model using a permutation procedure (42). No significant differences in mean coactivation, global anatomical coactivation nor community coactivation were observed across group comparisons (*p*’s>0.05; Table 1). However, when comparing coactivation of distinct anatomical partitions, female alcohol naive mice had greater hypothalamic and midbrain coactivation relative to male alcohol naive mice (*p*’s<0.05; Table 2). Female alcohol naive and binge drinking mice had increased hypothalamic coactivation compared to female alcohol anticipating counterparts (p’s<0.05; Table 2). Whilst male alcohol naive mice had increased forebrain coactivation compared to male alcohol anticipating and binge drinking counterparts (*p*’s<0.05; Table 2).

**Table 1.** Comparison of global network metrics between treatment groups, within and between sexes. \**p*<0.05, \*\**p*<0.01. ExtAmyg, extended amygdala; Hipp., hippocampus; Hypothal, hypothalamus.

| *Comparison* | *Mean coactivation* | *Anatomical modularity* | *Community modularity* |
| --- | --- | --- | --- |
| ♂ alcohol naïve v. | 0.9191 | 0.9008 | 0.5245 |
| ♂ alcohol anticipating |  |  |  |
| ♂ alcohol naïve v. | 0.3459 | 0.7198 | 0.3726 |
| ♂ binge drinking |  |  |  |
| ♂ alcohol anticipating v. | 0.5777 | 0.3544 | 0.4199 |
| ♂ binge drinking |  |  |  |
| ♀ alcohol naïve v. | 0.8343 | 0.2857 | 0.5814 |
| ♀ alcohol anticipating |  |  |  |
| ♀ alcohol naïve v. | 0.9578 | 0.7734 | 0.0948 |
| ♀ binge drinking |  |  |  |
| ♀ alcohol anticipating v. | 0.8009 | 0.2534 | 0.2308 |
| ♀ binge drinking |  |  |  |
| ♂ alcohol naïve v. | 0.6603 | 0.9983 | 0.1189 |
| ♀ alcohol naïve |  |  |  |
| ♂ alcohol anticipating v. | 0.4174 | 0.3281 | 0.3817 |
| ♀ alcohol anticipating |  |  |  |
| ♂ binge drinking v. | 0.8365 | 0.3750 | 0.2764 |
| ♀ binge drinking |  |  |  |

**Table 2.** Comparison of anatomical partition modularity between treatment groups, within and between sexes. \**p*<0.05, \*\**p*<0.01. ExtAmyg, extended amygdala; Hipp., hippocampus; Hypothal, hypothalamus.

| *Comparison* | *Forebra* | *ExtAmy* | *Hypothal* | *Thalamus* | *Hipp.* | *Midbrai* | *Hindbrai* |
| --- | --- | --- | --- | --- | --- | --- | --- |
| ♂ alcohol naïve v. | **0.0251\*** | 0.4528 | 0.3485 | 0.5052 | 0.3724 | 0.2599 | 0.8438 |
| ♂ alcohol anticipating |  |  |  |  |  |  |  |
| ♂ alcohol naïve v. | **0.0256\*** | 0.5181 | 0.0676 | 0.5647 | 0.6935 | 0.0769 | 0.3572 |
| ♂ binge drinking |  |  |  |  |  |  |  |
| ♂ alcohol anticipating v. | 0.6905 | 0.6418 | 0.3961 | 0.9113 | 0.3723 | 0.5319 | 0.3788 |
| ♂ binge drinking |  |  |  |  |  |  |  |
| ♀ alcohol naïve v. | 0.1410 | 0.8275 | **0.0058\*\*** | 0.2069 | 0.1224 | 0.5728 | 0.2279 |
| ♀ alcohol anticipating |  |  |  |  |  |  |  |
| ♀ alcohol naïve v. | 0.2844 | 0.5734 | 0.4411 | 0.9225 | 0.9761 | 0.1900 | 0.8928 |
| ♀ binge drinking |  |  |  |  |  |  |  |
| ♀ alcohol anticipating v. | 0.4351 | 0.5584 | **0.0238\*** | 0.2922 | 0.2078 | 0.1753 | 0.2771 |
| ♀ binge drinking |  |  |  |  |  |  |  |
| ♂ alcohol naïve v. | 0.2098 | 0.9458 | **0.0140\*** | 0.8351 | 0.1923 | **0.0058\*\*** | 0.8520 |
| ♀ alcohol naïve |  |  |  |  |  |  |  |
| ♂ alcohol anticipating v. | 0.0887 | 0.3636 | 0.3550 | 0.3831 | 0.4978 | 0.5087 | 0.2511 |
| ♀ alcohol anticipating |  |  |  |  |  |  |  |
| ♂ binge drinking v. | 0.4545 | 0.9307 | 0.6558 | 0.6710 | 0.3745 | 0.7186 | 0.3788 |
| ♀ binge drinking |  |  |  |  |  |  |  |

**Figure 1.**
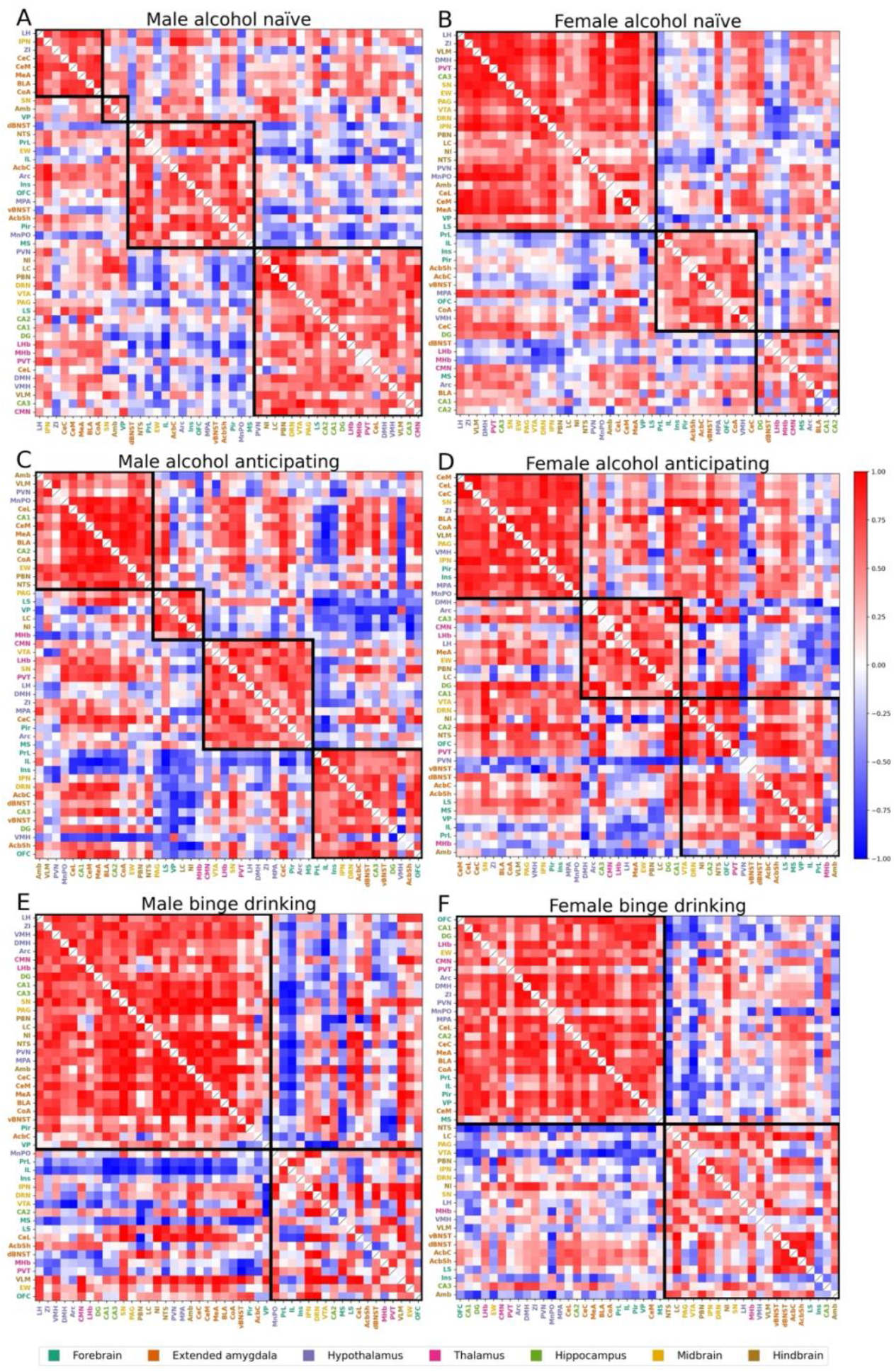
Louvain community consensus clustering of strongly correlated brain regions revealed differences in modularity between treatment groups, within and between sexes. Alcohol naive male **(A)** and female **(B)** mice had 4 and 3 community partitions respectively. For alcohol anticipating mice, males had 4 **(C)** and females had 3 **(D)** community partitions. Both male **(E)** and female **(F)** binge drinking mice had 2 community partitions. Abbreviations: ExtAmyg, extended amygdala; Hypothal, hypothalamus. Refer to Table S1 for brain region abbreviations.

We next assessed c-Fos expression of individual brain regions (see Table S5 for complete statistical analysis). Sex x treatment group interactions were found for the dBNST, BLA, LH, MHb and VTA. Bonferroni *post-hoc* analysis revealed reduced c-Fos expression in the VTA of male binge drinking (*p*=0.009) and alcohol anticipating mice (*p*=0.0002) relative to alcohol naïve counterparts, whilst VTA expression was also reduced in female compared to male alcohol naïve mice (*p*=0.006). There was greater LH c-Fos expression in alcohol anticipating females than alcohol naïve counterparts (*p*=0.006). c-Fos expression in the dBNST (*p*=0.022) and MHb (*p*=0.037) was greater in binge drinking compared to alcohol naïve females (Fig. S3).

Notably, the BLA was the only region where a difference in c-Fos expression was found between both the same-sex alcohol naïve group and same treatment group of the opposite sex. BLA c-Fos expression was greater in female binge drinking (*p*=0.003) and alcohol anticipating mice (*p*=0.002) than alcohol naïve counterparts, and in female compared to male binge drinking mice (*p*=0.047; Fig. 2A), suggesting increased BLA engagement following binge drinking in female mice (Fig. 2B; see Table 3 and Fig. S3 for main effect differences).

**Table 3.**
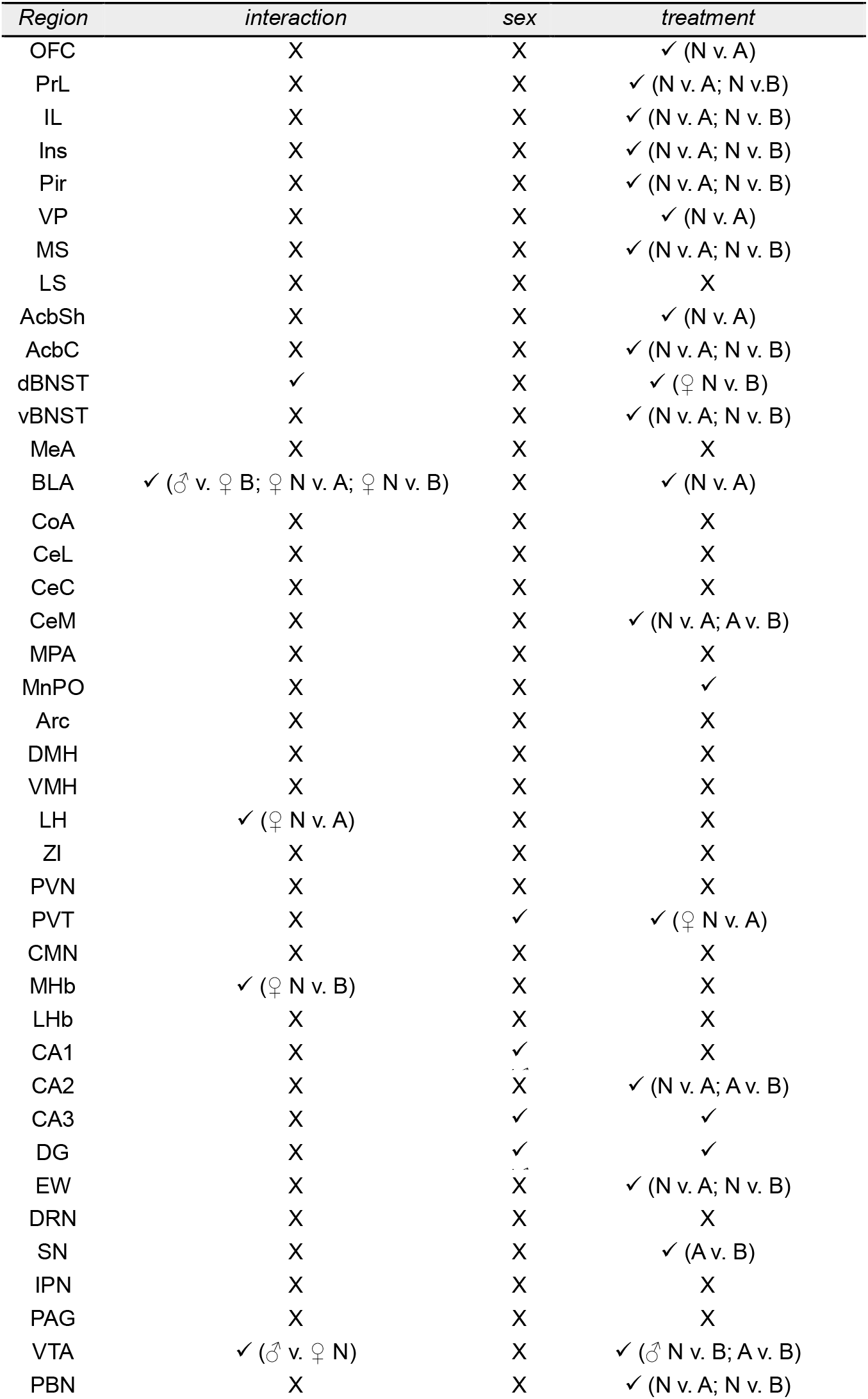
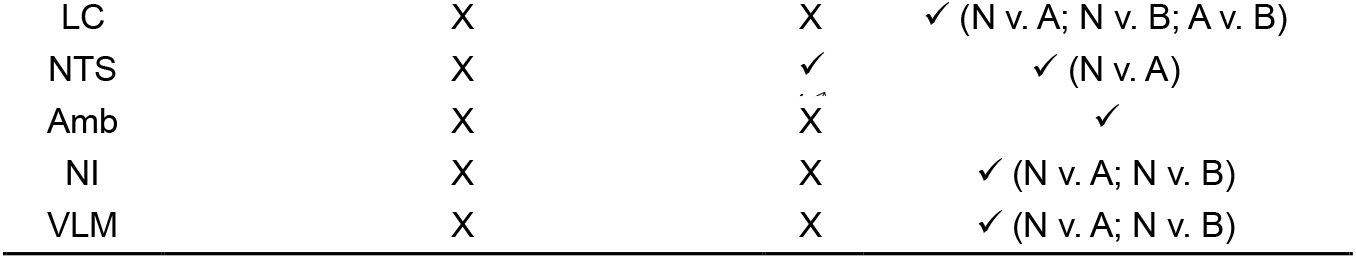
Summary of interactions and main effects of c-Fos expression analyses. A, alcohol anticipating; B, binge drinking; N, alcohol naïve. Please refer to Table S1 for brain region acronyms.

**Figure 2.**
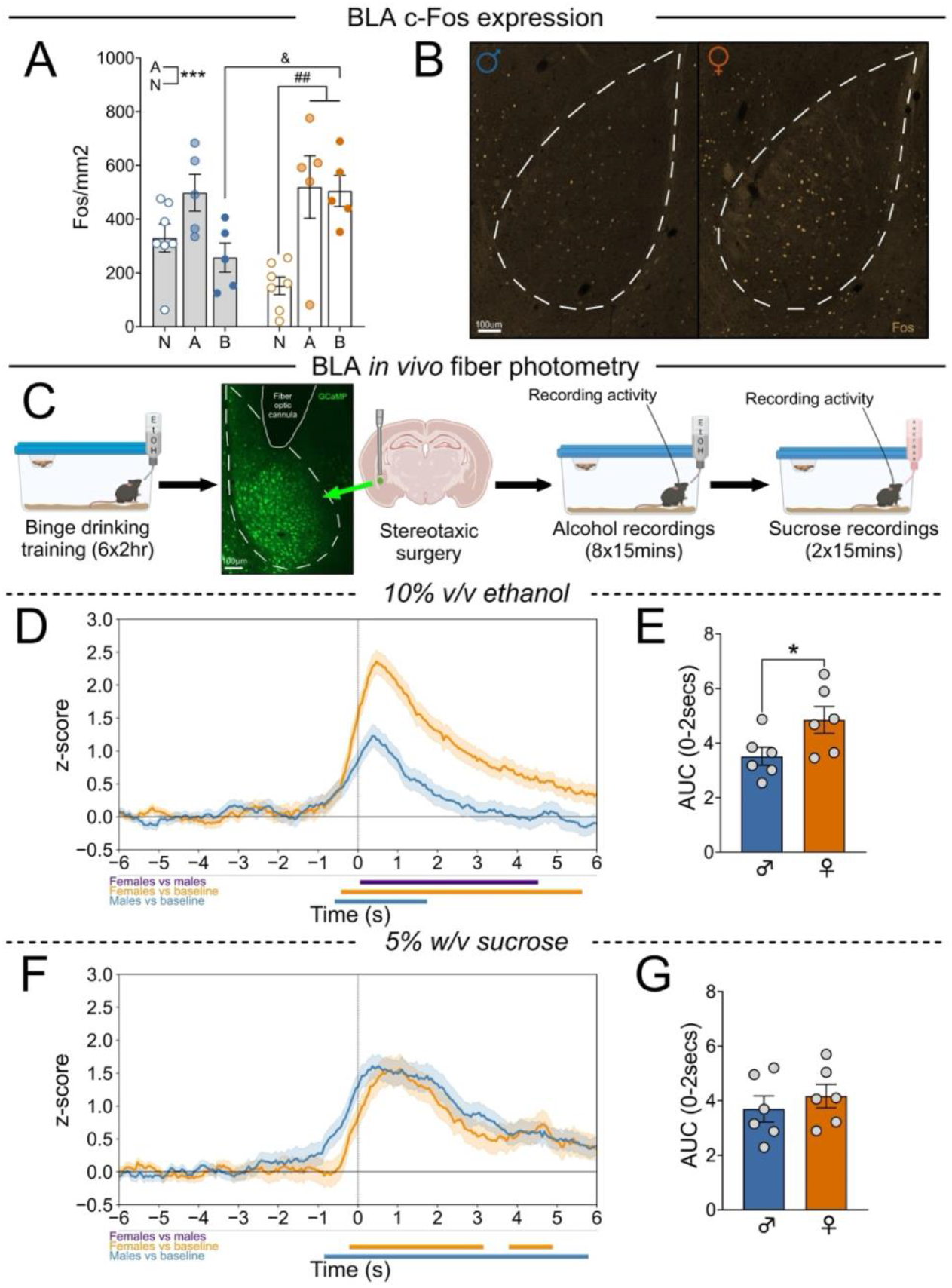
BLA activation, ex vivo and in vivo, in response to alcohol is greater in female compared to male mice. **(A)** Female binge drinking mice had increased Fos/mm2 expression compared to male binge drinking and female naïve mice (n=5/7/sex/group). **(B)** Representative images of Fos-protein expression in the BLA of a male (left) and female (right) mouse following binge drinking. **(C)** Timeline of fiber photometry experimental procedure with surgical intervention and representative image of viral injection site and probe placement in the BLA. **(D)** BLA activity in male (blue line below graph) and female (orange line below graph) was increased in response to onset of alcohol drinking (vertical line), with this response being greater and more prolonged in females (purple line below graph; n=6/sex). **(E)** Average AUC 0-2secs from onset of alcohol drinking was greater in female compared to male mice. **(F)** Onset of sucrose drinking increased BLA activity in male (blue line below graph) and female (orange line below graph) mice, with this response being comparable between sexes over the timecourse. **(G)** There was no difference in average AUC 0-2secs from onset of sucrose drinking between sexes. Data presented as mean ± SEM. *p<0.05, ^ = difference between treatment groups (^^^p<0.001), & = difference between sexes for the same treatment group (&p<0.05), # = difference between treatment groups within sex (##p<0.01). Bars at bottom of **(D)** and **(F)** were determined via bootstrapped confidence intervals (99%). Abbreviations: A, anticipating; AUC, area under the curve; B, binge drinking; N, naïve; s, seconds.

We subsequently used real-time *in vivo* fiber photometry to further probe the temporal nature of alcohol-induced BLA activity (Fig. 2C). Both sexes showed increased BLA activity at the onset of alcohol intake relative to baseline activity; however, this response was significantly stronger and more sustained in female compared to male mice (99% CI, consecutive threshold of 0.5s; Fig. 2D) with the average area under the curve (AUC) greater in females (*p*<0.05; Fig. 2E). Examination of response to sucrose showed increases in BLA activity at the onset of consumption (99% CI, consecutive threshold of 0.5s; Fig. 2F), effects that were comparable between sexes as shown by AUC analysis (*p*>0.05; Fig. 2G; see Table S6 for complete statistical analysis).

### Global BLA inhibition reductions in reward intake in both sexes

Given we observed enhanced BLA activity to alcohol intake in female mice, we next sought to functionally test the role of the BLA in binge drinking through bilateral chemogenetic inhibition (Fig. 3A). CNO administration did not alter cumulative nor total alcohol intake in male (Fig. 3B) or female control mice (*p*’s>0.05; Fig. 3C). Bilateral BLA inhibition reduced cumulative alcohol intake in male mice (main effect of treatment, *p*<0.05; Fig. 3D) at the 2hr (*p*=0.0007), 3hr (*p*=0.0002) and 4hr timepoints (*p*=0.002), with a trend toward reduced total intake (*p*=0.057; Fig. 3D). In females, BLA inhibition also reduced cumulative intake (main effect of treatment, *p*<0.05; Fig. 3E) at the 3hr (*p*=0.0008) and 4hr timepoints (*p*=0.0004), and total alcohol intake (*p*<0.05; Fig. 3E). Assessment of delta (Δ) alcohol intake following CNO relative to vehicle administration, revealed a significant reduction in hM4Di mice compared to controls (main effect of viral group, *p*<0.05), while no difference between sexes was observed (main effect of sex, *p*>0.05; Fig. 3F). Viral spread shown in Fig. 3G-H.

**Figure 3.**
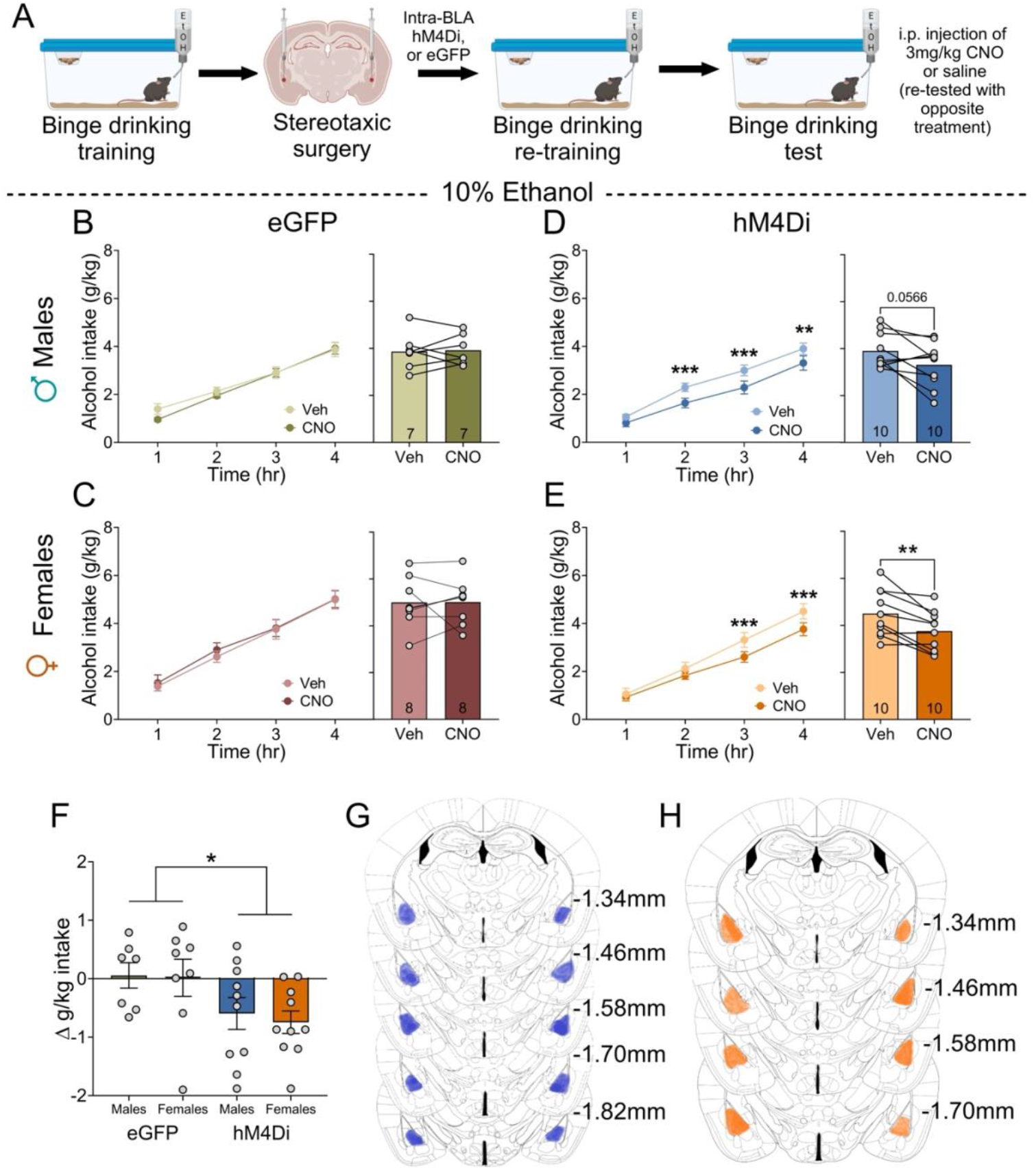
Chemogenetic BLA inhibition reduces alcohol intake in both sexes. **(A)** Schematic representation of experimental timeline with surgical intervention. CNO administration did not alter cumulative (left) nor total (right) alcohol intake in male **(B)** and female **(C)** mice eGFP control virus (n=7-8/sex). **(D)** BLA inhibition reduced cumulative intake in male mice (n=10) at the 2hr, 3hr and 4hr timepoints (left) and trended toward reducing total intake (right). While, in female mice **(E)**, this reduced cumulative intake at the 3hr and 4hr timepoints (left) and total alcohol intake (right; n=10). **(F)** Δ alcohol intake following CNO administration was reduced in hM4Di mice relative to eGFP control counterparts. Visual representation of viral targeting of the BLA in male **(G)** and female **(H)** mice. Data presented as mean ± SEM. **p<0.01, ***p<0.001. Abbreviations: BLA, basolateral amygdala; CNO, clozapine-N-oxide; g, gram; hr, hour; kg, kilogram; veh, vehicle.

We next assessed whether these reductions in alcohol intake generalized to natural reward by comparing sucrose consumption. Again, CNO had no effect on cumulative or total sucrose intake in male (Fig. S3A) or female (Fig. S3B) control mice (*p*’s>0.05). In males, bilateral BLA inhibition reduced cumulative intake (main effect of treatment, *p*<0.05; Fig. S3C) at the 2hr (*p*=0.0002), 3hr (*p*<0.0001) and 4hr (*p*<0.0001) timepoints, and total sucrose intake (*p*<0.05; Fig. S3C). Similarly, in females, bilateral BLA inhibition reduced cumulative intake (main effect of treatment, *p*<0.05; Fig. S3D) at the 3hr (*p*=0.003) and 4hr (*p*<0.0001) timepoints, and total sucrose intake (*p<*0.05; Fig. S3D). No significant difference in Δ sucrose intake following CNO administration was observed between hM4Di and control mice or between sexes (main effects of viral group and sex, *p*’s>0.05; Fig. S3E; see Table S7 for complete statistical analysis).

### c-Fos activation of the BLA→AcbC projection is greater in female compared to male binge drinking mice

Given the BLA is a key central processing hub with several distinct and defined projections (43-46), we next assessed potential sex differences in the engagement of discrete BLA efferent projections during binge drinking using CTβ and c-Fos dual labelling (Fig. 4A). Consistent with our initial findings, there was increased BLA c-Fos expression in female compared to male binge drinking mice (*p*<0.05; Fig. 4B). Notably, for the BLA→AcbC projection (Fig. 4C), females had increased CTβ expression and %CTβ-positive c-Fos cells (*p*’s<0.05; Fig. 4D) relative to males. For the BLA→mPFC projection (Fig. 4E), we observed reduced CTβ expression and %CTβ-positive c-Fos cells (*p*’s<0.05; Fig. 4F) in female compared to male binge drinking mice. No sex differences in CTβ expression or %CTβ-positive Fos c-cells were observed for the BLA to AcbSh (Fig. 4G-H), BNST (Fig. 4I-J) and vHipp (Fig. 4K-L) projections (*p*’s>0.05; see Table S8 for complete statistical analysis).

**Figure 4.**
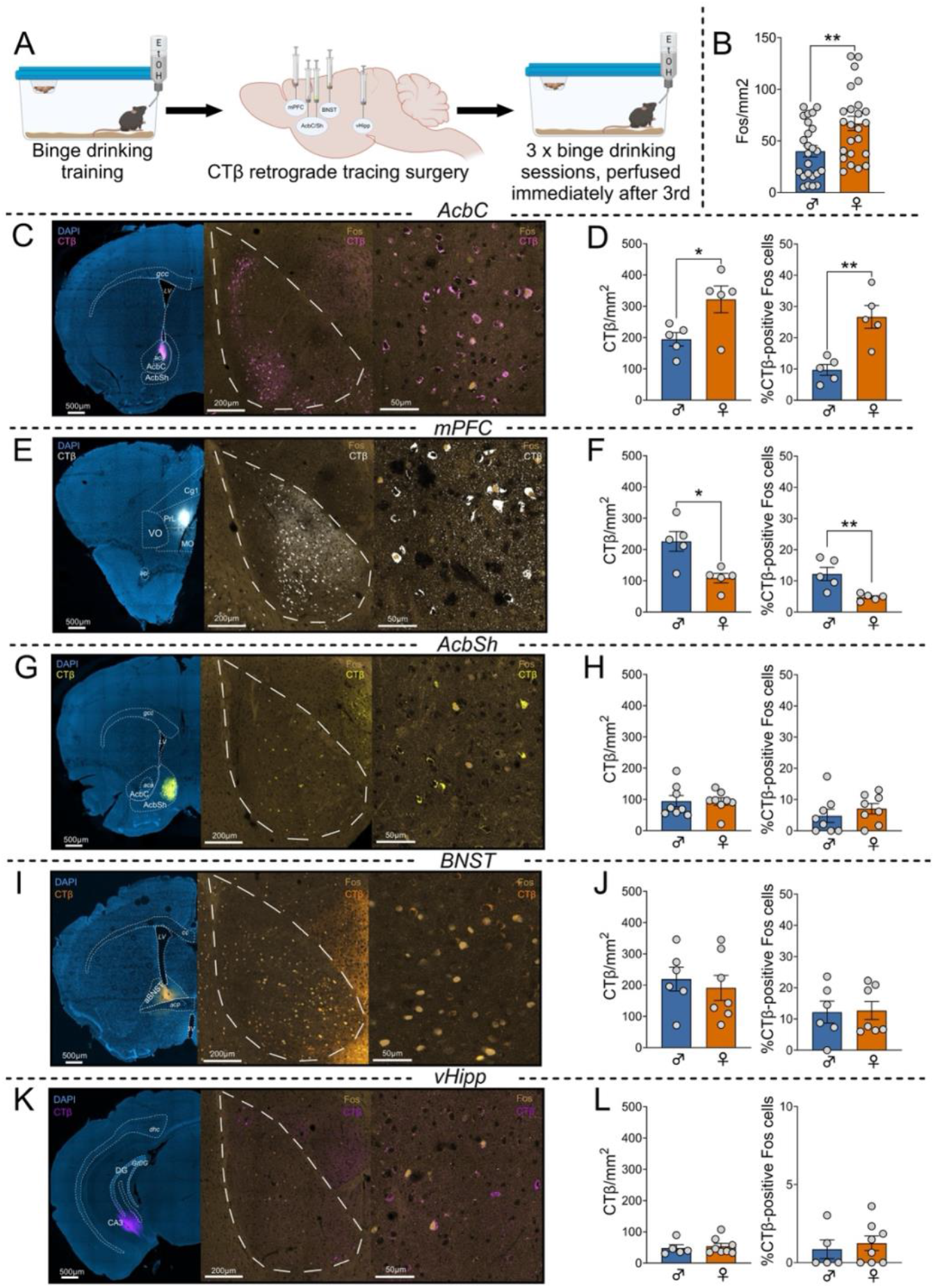
Activation of the BLA→AcbC projection is greater in female compared to male binge drinking mice. **(A)** Schematic representation of experimental timeline with surgical intervention. **(B)** Females had increased c-Fos/mm2 expression in the BLA compared to males following binge drinking (n=23/sex). **(C)** Representative images of a mPFC CTβ injection site (left), and overview (middle) and higher magnification (right) of BLA CTβ and c-Fos expression. **(D)** CTβ/mm2 expression (left) and percentage of CTβ-positive Fos cells (right) for the BLA→mPFC projection is greater in male compared to female mice following binge drinking (n=5/sex). **(E)** Representative images of an AcbC CTβ injection site (left), and overview (middle) and higher magnification (right) of BLA CTβ and c-Fos expression. **(F)** Increased CTβ/mm2 expression (left) and percentage of CTβ-positive Fos cells (right) was found for the BLA→AcbC projection in female relative to male binge drinking mice (n=5/sex). **(G)** Representative images of an AcbSh CTβ injection site (left), and overview (middle) and higher magnification (right) of BLA CTβ and c-Fos expression. **(H)** No sex difference in CTβ/mm2 expression (left) and percentage of CTβ-positive Fos cells (right) was found for the BLA→AcbSh projection (n=8/sex). **(I)** Representative images of a BNST CTβ injection site (left), and overview (middle) and higher magnification (right) of BLA CTβ and c-Fos expression. **(J)** CTβ/mm2 expression (left) and percentage of CTβ-positive Fos cells (right) for the BLA→BNST projection were similar between sexes (n=6-7/sex). **(K)** Representative images of a vHipp CTβ injection site (left), and overview (middle) and higher magnification (right) of BLA CTβ and c-Fos expression. **(L)** Male and female mice had comparable CTβ/mm2 expression (left) and percentage of CTβ-positive Fos cells (right) for the BLA→vHipp projection projection (n=5 males/8 females). Data presented as mean ± SEM. *p<0.05, **p<0.01. Abbreviations: aBNST, bed nucleus of the stria terminalis; aca, anterior commissure anterior part; AcbC, nucleus accumbens core; AcbSh, nucleus accumbens shell; aci, anterior commissure intrabulbar part; acp, anterior commissure posterior; BLA, basolateral amygdala; BNST, bed nucleus of the stria terminalis; cc, corpus callosum; Cg1, cingulate cortex area 1; CTβ, Cholera Toxin Subunit B; DG, dentate gyrus; dhc, dorsal hippocampal commissure; gcc, genu of the corpus callosum; GrDG, granular layer of the dentate gyrus; IL, infralimbic cortex; LV, lateral ventricle; MO, medial orbital cortex; mPFC, medial prefrontal cortex; PrL; prelimbic cortex; vHipp, ventral hippocampus; VO, ventral orbital cortex.

### BLA→AcbC pathway inhibition reduces binge alcohol intake in female but not male mice

To determine whether the BLA→AcbC projection mediates binge drinking specifically in female mice, we used pathway-specific chemogenetics to inhibit this projection (Fig. 5A). BLA→AcbC pathway inhibition did not alter cumulative (main effect of treatment) nor total alcohol intake in male mice (*p’*s>0.05; Fig. 5B). Bilateral inhibition in females resulted in a significant treatment x time interaction (*p<*0.05) and trend toward a main effect of treatment (*p*=0.059), with Bonferroni *post-hoc* analysis revealing significant intake reductions at the 3hr (*p*=0.0003) and 4hr timepoints (*p*<0.0001; Fig. 5C). Total alcohol intake in female mice was also reduced (*p*<0.05; Fig. 5C). Further, there was a sex difference in Δ alcohol intake following CNO administration relative to vehicle, with females having reduced Δ alcohol intake compared to male mice (*p*<0.05; Fig. 5F).

**Figure 5.**
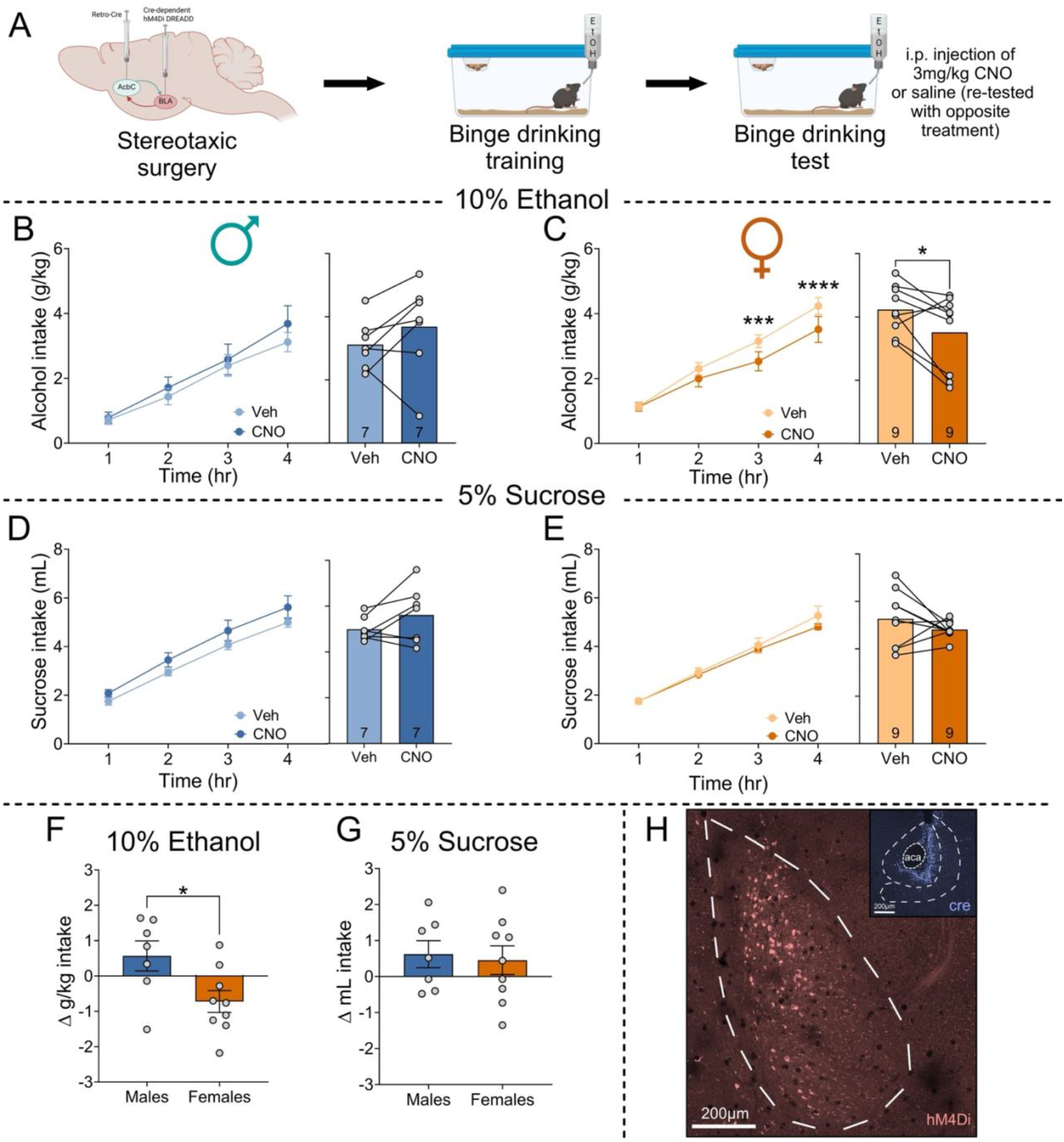
BLA→AcbC pathway-specific inhibition reduces female binge drinking in a sex- and alcohol-specific manner. **(A)** Schematic representation of experimental timeline with surgical intervention. **(B)** Inhibition of the BLA→AcbC projection did not alter cumulative (left) nor total (right) alcohol intake in male mice (n=7). **(C)** BLA→AcbC pathway inhibition reduced cumulative alcohol intake at the 3hr and 4hr timepoints (left) and total alcohol intake (right) in female mice (n=9). BLA→AcbC pathway-specific inhibition did not alter cumulative (left) nor total (right) sucrose intake in male **(D)** and female **(E)** mice. **(F)** CNO administration reduced Δ alcohol intake specifically in female mice, while not altering Δ sucrose intake in either sex **(G). (H)** Representative images of cre-dependent DREADD expression in the BLA and targeting of the AcbC with the retrograde Cre-carrying virus. Data presented as mean ± SEM. *p<0.05, ***p<0.001, ****p<0.0001. Abbreviations: AcbC, nucleus accumbens core; BLA, basolateral amygdala, CNO, clozapine-N-oxide; g, gram; hr, hour; kg, kilogram; mg, milligram.

We next assessed whether this reduction in intake in female mice was specific to alcohol, or also reduced sucrose intake. Inhibition of the BLA→AcbC projection did not alter cumulative nor total sucrose intake in male (Fig. 5D) or female (Fig. 5E) mice (*p*’s<0.05). There was no sex difference in Δ sucrose intake following CNO administration (*p*>0.05; Fig. 5G). Representative images of injection sites shown in Fig. 5H (see Table S9 for complete statistical analysis).

## Discussion

We identify brain region and network level sex differences across alcohol-related states, revealing heightened BLA activation in response to alcohol intake in female compared to male mice. This increased engagement was, in part, associated with preferential activation of the BLA→AcbC projection, with functional inhibition of this pathway sex-specifically reducing binge alcohol intake in female mice. Together, these data underscore the nuanced, sex-specific neurobiological drivers of alcohol use, and identify the BLA→AcbC projection as a pathway that contributes to excessive alcohol consumption in a sex-specific manner.

We first examined network level differences across sexes and treatment to characterize coordinated patterns of brain activation underlying alcohol-related behaviors (41, 47, 48). Louvain community detection and consensus clustering of regions with similar patterns of activation into communities/modules revealed male and female mice to exhibit fewer modules than naïve and alcohol anticipating counterparts, suggesting reduced network modularity following excessive alcohol intake. Consistent with this, binge drinking and ethanol dependence have previously been reported to reduce modularity in male mice (28, 42, 49). However, this appears sensitive to experimental context, as other studies have reported increased modularity following a 4hr DID session in male mice relative to naïve counterparts (50). In contrast to our findings, female mice with a frontloading drinking phenotype (28) and following binge drinking (50, 51) also showed greater modularity compared to naïve controls. This suggests network modularity is dependent on experimental parameters, including drinking model, duration of ethanol exposure, euthanasia timepoint, and tissue processing methodology.

Whilst examining region-specific c-Fos expression, we found both alcohol state related differences across several nodes (dBNST, LH, BLA, MHb VTA) and sex-dependent effects within alcohol naïve (CA1, VTA), alcohol anticipating (NTS) and binge drinking (DG, BLA) groups. Notably, the BLA was the only region to show a sex difference in c-Fos within binge drinking mice, and an increase in c-Fos relative to female naïve controls. In line with this, *in vivo* fiber photometry revealed enhanced and prolonged BLA activation at the onset of alcohol, but not sucrose, drinking in females, supporting a role for the BLA in female alcohol consumption, consistent with prior reports (50, 52). Mechanistically, acute binge-like alcohol concentrations enhance inhibitory tone via activation of BLA GABAergic neurons (∼15%) (53) that regulate glutamatergic neurons (∼85%) (53) in a sex-dependent manner in male rats (54). This raises the possibility that stronger inhibitory tone may constrain BLA glutamatergic activation in males, relative to females, in response to high levels of alcohol exposure. Importantly, these sex differences appear specific to acute binge-like exposure, as to date, no sex differences have been reported following chronic drinking (55).

Using chemogenetics we global BLA inhibition led to reduced reward intake (alcohol and sucrose) in both sexes, indicating a generalized disruption of reward behavior. Given the established roles for the BLA in reward processing and motivated behavior (56-58), such a broad reduction in intake is not unexpected; indeed, bilateral BLA lesions reduce sucrose intake in male rats (59). Given the intricate microcircuits within the BLA, this inhibition likely caused broad disruption in signaling dynamics between interneuron and projection neurons (60, 61), thereby interfering with regulation of BLA outputs (61, 62) and downstream behavioral control.

As the BLA is a critical central processing hub with multiple anatomically and functionally distinct efferent projections implicated in reward, motivation and anxiety (43-46), we assessed involvement of discrete BLA efferent projections. No sex differences were observed in the activation, or density, of BLA projections to the AcbSh, BNST and vHipp. Previous work has demonstrated a role for the BLA→vHipp pathway in appetitive reward seeking in male rats (63), and specific BLA→Acb and BLA→PFC neuronal populations in the formation and expression of alcohol-cue associations in male mice (64). However, whether these generalize across the sexes remains unclear. Evidence from chronic intermittent alcohol exposure and withdrawal (CIE/WD) models indicate that sex differences in BLA projection function may be both behavior and exposure dependent (65). Consistent with this, we observed an increased percentage of c-Fos in BLA→mPFC projecting cells in binge drinking male mice compared to females. The biological significance of this finding should be considered with caution given males had both increased CTβ and reduced c-Fos expression relative to females, which may bias proportional co-expression estimates.

Notably, we identified increased engagement of the BLA→AcbC projection in female compared to male binge drinking mice, revealing a potential sex-specific pathway underlying binge alcohol intake. Consistent with this, chemogenetic inhibition of the BLA→AcbC pathway selectively reduced alcohol consumption in females, but not males, supporting the functional relevance of this projection in female binge drinking. While the effect size was modest, this suggests this pathway represents one of several circuits contributing to binge drinking in female mice (30, 31). A recent study observed no sex differences in the engagement of BLA→Acb projections following binge alcohol drinking (29); however, specificity of targeting may account for the differences in findings given we observed no sex differences in BLA→AcbSh projection activation. The BLA→AcbC pathway specifically mediates the conditioned reinforcement and cue-driven drug seeking for psychostimulants (66, 67), although these studies were both conducted in male rodents only. The majority of BLA→AcbC projections are glutamatergic (>90%) (68) and interestingly, studies have reported sex differences in both structural synaptic connectivity and glutamatergic and dopaminergic signaling in the AcbC (69-72). Notably, female rodents display a higher AcbC AMPA/NMDA ratio and larger readily releasable pool of glutamate in the AcbC (72), which has been posited to be projection-specific (72), but not attributable to vHipp inputs (73). Together, these sex-dependent synaptic features provide a plausible substrate through which heightened recruitment of the BLA→AcbC pathway may preferentially drive binge drinking in females.

In summary, our work suggests that binge alcohol consumption engages both overlapping and distinct neural circuits in male and female mice. By interrogating network level analyses, regional activity mapping, and pathway-specific manipulations, our findings identify a sex-dependent circuit projection from the BLA to AcbC that contributes to excessive alcohol consumption in females. Together, our data highlight the importance of incorporating sex as a biological variable in systems neuroscience approaches, supporting further interrogation of sex informed treatment approaches for AUD.

## Acknowledgements, funding and author contributions

We thank the Florey Core Animal Services and Florey Microscopy Facility for their assistance. This project was supported by a University of Melbourne Early Career Researcher grant (XJM), National Health and Medical Research Council (NHMRC) Ideas grant (2002830, LCW), NHMRC Emerging Leader Fellowship (2008344, LCW). AJL is supported by an NHMRC Synergy Grant (2009851). XM, LTU, KLH were supported by Australian Research Training Program Scholarships and HD by an Australian Rotary Health/Rob Henry Memorial PhD scholarship. EJC was supported by the Australian Research Council (DE230100401). We acknowledge support from the Victorian State Government Operational Infrastructure Scheme.

XJM Conceptualization; Data curation; Formal analysis; Funding acquisition; Investigation; Methodology; Supervision; Writing – original draft; Writing – review & editing. AJP Investigation; Methodology; Writing – review & editing. QTT Investigation; Methodology; Writing – review & editing. HD Formal analysis; Methodology; Writing – review & editing. LTU Investigation; Writing – review & editing. KLH Investigation; Writing – review & editing. BKR Investigation; Methodology; Writing – review & editing. RGA Investigation; Writing - review & editing. EJC; Writing – review & editing. AJL; Resources; Supervision; Writing – review & editing. RMB Conceptualization; Writing – review & editing. LCW Conceptualization; Data curation; Formal analysis; Funding acquisition; Investigation; Methodology; Project administration; Supervision; Writing – original draft; Writing – review & editing.

## Disclosures

The authors report no potential conflicts of interest.

## Notes

### Competing Interest Statement

The authors have declared no competing interest.

