## Supplementary material for "Sex differences in neural circuits driving binge drinking: A female-specific role for an amygdalo-striatal pathway"

**Supplementary Materials & Methods**

*Animals*

C57BL6J mice (n=149) were acquired from the Australian Resource Centre (ARC, Perth, Australia). Mice were single-housed and maintained in a temperature-controlled room (21°C) under a 12hr light/dark cycle (lights off at 0700) with *ad libitum* access to food (laboratory chow, Barastoc) and water, except where detailed. All mice were acclimated to the experimental holding room for one week prior to the commencement of experimentation (~8 weeks of age). All experiments were performed in accordance with the Prevention of Cruelty to Animals Act (2004), following the guidelines of the National Health and Medical Research Council (NHMRC) Australian Code of Practice for the Care and Use of Animals for Experimental Purposes (2013), and approved by the Florey Institute of Neuroscience and Mental Health Animal Ethics Committee.

*Binge drinking procedure*

Binge drinking sessions consisted of the home-cage water bottle being replaced with a bottle containing 10% v/v ethanol or 5% w/v sucrose (diluted in tap water) 3 times/week (Monday, Wednesday, Friday), 3hrs into the dark phase for 2hrs (1, 2). Mice had free access to the solution during this time, with the water bottle returned immediately following the 2hr session. A bottle of 10% v/v ethanol or 5% w/v sucrose was also placed into an empty cage during each session and weighed at the same timepoints to account for potential spillage. Once drinking levels stabilized, mice underwent either a final 2hr binge drinking session or an extended 4hr binge drinking test session.

The extended 4hr binge drinking test session involved mice receiving an intraperitoneal (i.p.) injection of vehicle (saline; 10mL/kg) or CNO (3mg/kg; Sigma-Aldrich), 40min prior to the presentation of the 10% v/v ethanol or 5% w/v sucrose bottle. The solution bottle was weighed each hour across the 4hr test to track the time course of intake. Mice were then re-trained to baseline drinking levels before being tested with the opposite treatment condition (~1 week between sessions; counterbalanced).

*g/kg intake calculation*

Total intake was calculated using the volume consumed in milliliters (mL), subtracting the spillage volume and multiplying by the density of 10% v/v ethanol. The grams per kilogram (g/kg) was calculated as follows: total alcohol consumed (grams) x alcohol concentration (0.1) * density of alcohol (0.79)/(weight of mouse (grams)/1000).

*Stereotaxic surgery*

Mice were anaesthetized with isoflurane (5% induction; 2% maintenance) and placed in a stereotaxic frame (Stoelting Co., Illinois, USA). The surgical site was shaved and cleaned with betadine, and a small incision was made along the midline to expose the skull.

For *in vivo* fiber photometry, a burr hole was drilled unilaterally followed by microinjection of GCaMP6m (AAV9-Syn-GCaMP6m-WPRE-SV40, 250nL, 1/3 dilution in PBS, 100838, Addgene) into the BLA (AP: -1.80mm; ML: +/- 3.20mm; DV: -5.00mm). A 400µm fiber optic cannula (Neurophotometrics) was implanted above the BLA (AP: -1.80mm; ML: +/- 3.20mm; DV: -4.85mm) and anchored to the skull using dental cement (Vertex-Dental) and screws.

For chemogenetic inhibition of the BLA, burr holes were drilled and mice were bilaterally microinjected with either an inhibitory DREADD (AA5-hSyn-hM4Di(Gi)-mCherry, 80nL, titer: 1.8 x $10^{13}$ Gc/mL, Lot v169738, 50465, Addgene) or control (AAV5-hSyn-eGFP, 80nL, 2.26 x $10^{13}$ Gc/mL, Lot v128149, 50465, Addgene) virus into the BLA (AP: -1.80mm; ML: +/- 3.20mm; DV: -5.00mm).

To investigate activation of BLA efferent projections after binge drinking, burr holes were drilled above a target region(s) with 40nL of one or two of the following cholera toxin subunit β (CTβ) conjugates CTβ-488 (Cat# C34775), CTβ-555 (Cat# C34776) and/or CTβ-647 (Cat# C34778; 2%, Invitrogen, Carlsbad, CA, USA) unilaterally microinjected into the mPFC (AP: +1.70mm; ML: +/-0.40mm; DV: -2.60mm), AcbC (AP: +1.10mm; ML: +/-0.85mm; DV: -4.60mm), AcbSh (AP: +1.10mm; ML: +/-0.55mm; DV: -4.60mm), BNST (AP: +0.10mm; ML: +/-1.00mm; DV: -4.00mm) and/or vHipp (AP: -3.20mm; ML: +/-3.30mm; DV: -4.25mm).

For pathway specific inhibition of the BLA→AcbC projection, burr holes were drilled bilaterally above the AcbC and BLA, with 150nL of a retrograde Cre-carrying virus (#55637-AAVrg, Addgene) and a Cre-dependent inhibitory DREADD (AAV-hSyn-DIO-hM4Di(Gi)-mCherry; Lot v143762, #44362-AAV2, Addgene) being microinjected into the AcbC (AP: +1.10mm; ML: +/-0.85mm; DV: -4.60mm) and BLA (AP: -1.80mm; ML: +/- 3.20mm; DV: -5.00mm) respectively.

All microinjections were delivered at a rate of 1nL/sec using a Nanoliter Microinjection Pump (RWD Life Science, Shenzhen, China), with the micropipette remaining in place for an additional 5mins to minimize spread and backtracking. Mice were administered with meloxicam (3mg/kg$^{-1}$ s.c.), the antibiotic baytril (3mg/kg$^{-1}$, s.c.) and saline (10mL/kg$^{-1}$, s.c.) to facilitate recovery. Mice were given at least one week to recover from surgery prior to recommencing behavioral training.

*In vivo fiber photometry*

A Neurophotometrics fiber photometry system (FP3002; Neurophotometrics, MBF, Bioscience, USA) was used for fiber photometry recordings, combined with Bonsai (open-source, V 2.8.5) for system control. Patch cables (0.37NA, 400μm core; Doric) transmitted alternating 470nm (GCaMP) and 415nm (isosbestic) LEDs at a 40Hz sampling rate (20Hz per channel). Raw fluorescence signals obtained were processed using the following open-source GUI available on GitHub (https://github.com/H-Dempsey/Fiber_photometry_analysis_NPM).

For fiber photometry recording sessions, mice were habituated to the behavior room for at least 30mins prior to the commencement of recordings. Each session consisted of a 15min recording period, the first 2mins being a baseline period where no solution was presented. Following this, either a 10% v/v ethanol bottle (first 8 sessions) or 5% w/v sucrose bottle (final 2 sessions) was presented for the remainder of the session.

*Transcardial perfusions*

Mice were anesthetized with pentobarbitone (80mg/kg; i.p.) and transcardially perfused with 12mL of phosphate buffered saline (PBS; 0.1M, pH 7.4), followed by 25mL of 4% w/v paraformaldehyde (PFA; Sigma-Aldrich) in PBS. Brains were extracted and post-fixed in 4% w/v PFA for one-hour (c-Fos immunohistochemistry) or overnight (viral targeting site validations), and then transferred into 30% w/v sucrose in PBS for ~24hr at 4°C. Brains were sectioned coronally (40μm) with a Leica Microsystems Cryostat, and stored in PBS containing 0.1% w/v sodium azide (Sigma-Aldrich) at 4°C until processed.

*Injection site validations*

To validate CTβ microinjection placements, sections of the targeted regions were washed in PBS (5mins), stained in PBS containing 0.5μL/mL of DAPI (Cat# 5748, Tocris, Bristol, UK; 5mins), rewashed in PBS (5mins) before being mounted on microscope slides and coverslipped with fluorescent mounting medium (DAKO, Carpentaria, CA, USA). For viral targeting validations, tissue was mounted directly onto microscope slides and coverslipped with fluorescent mounting medium (DAKO, Carpentaria, CA, USA).

*Immunohistochemistry*

A one-in-four series of the entire brain of all mice from the alcohol naïve, alcohol anticipating and binge drinking groups, or BLA for co-labelling with CTβ expression, were processed for c-Fos immunoreactivity. Free-floating sections were washed in 0.1M PBS (3 x 5mins) then pre-blocked with 10% normal donkey serum (NDS) and 0.5% v/v Triton-X100 (T-X) in PBS at room temperature (RT) for 2hr. Sections were then incubated in a primary antibody solution containing either goat anti-Fos (1:500, Santa Cruz Biotechnology, #SC-52-G; Fos mapping) or rabbit anti-Fos (1:1000; Cell Signaling Technology, #5348, Ser32, D82C12, Lot 4; Retrograde tracing), 2% NDS, 0.5% T-X in PBS for 24hrs at RT. Following this, sections were washed in PBS (3 x 5mins) and incubated at RT for 2hr in a secondary antibody solution of donkey anti-goat Alexa-fluor 488 (1:400, Life Technologies, A-21447; Fos mapping) or, for retrograde tracing co-labelling, donkey anti-rabbit Alexa-fluor 488 (1:400, Invitrogen, A21206) or 647 (1:400, Invitrogen, A-31573), 2% NDS and 0.5% T-X in PBS. Sections were washed in PBS (3 x 5mins), then mounted on microscope slides and coverslipped with fluorescence mounting medium (DAKO, Carpentaria, CA, USA). CTβ fluorescence was visualized endogenously, without amplification.

*Image acquisition and quantification*

A LSM780 Zeiss Axio Imager 2 confocal laser scanning microscope (Carl Zeiss AG, Jena, Germany) was used to digitally capture 20x magnification overview images of brain sections of alcohol naïve, alcohol anticipating and binge drinking mice. A LSM900 Zeiss Axio Imager 2 confocal laser scanning microscope was used to digitally capture 20x magnification images of c-Fos and CTβ expression in BLA sections and 10x magnification overview images of microinjection sites. For each experiment, the same laser settings were used for all mice. Quantification was performed using ImageJ software (National Institute of Health) by an experimenter blinded to the experimental conditions, with at least 3 sections of a given brain region required for an animal to be included in the analysis. Brain region identification was based on coordinates indicated in 'The Mouse Brain in Stereotaxic Coordinates Atlas' (3).

For mapping of c-Fos expression of alcohol naïve, alcohol anticipating and binge drinking mice, quantification encompassed 40 brain regions and their subdivisions (see Table S1), selected due to being previously implicated in reward and/or known to express sex steroid hormone receptors.

*Network level analyses*

To perform brain network analyses, the Python implementation (https://github.com/aestrivex/bctpy) of the Brain Connectivity Toolbox was used (4). Firstly, we constructed undirected, weighted, and signed Pearson inter-regional correlation matrices for each group (5-10). We then computed the global mean coactivation (mean *r* of each matrix), mean coactivation for each anatomical partition (based on the Allen Brain Atlas), and signed modularity according to anatomical partitions. To identify clusters of regions with similar patterns of activation in each group, we performed Louvain community detection (11-13) 1000 times and used the negative asymmetric objective function recommended in (13). From this, a weighted agreement matrix was constructed and consensus clustering performed with a tau of 0.5 to identify final community partitions (14). Given the order of detected Louvain communities is arbitrary, we ordered the correlations of each community according to decreasing mean coactivation.

We then compared mean coactivation, anatomical modularity (global and within anatomical divisions) and consensus clustering between treatment groups within sex, and in the same treatment group across sex through permutation tests (15). To so, we shuffled the group labels within each pairwise comparison and recomputed the Pearson correlation matrix, mean coactivation, signed anatomical modularity and signed modularity using the empirical consensus partitions, repeating this for every combination of pairwise group label shuffling.

The p-value was then computed using the following equation: $p_{stat} = \frac{\#\left\{\left|stat_{null}^{(i)}\right| \geq \left|stat_{emp}\right|\right\}}{N_{shufflings}}$

Our GitHub repo provides the input data and python script that were used to generate the figures and results reported (https://github.com/H-Dempsey/Brain_Network_Analysis_X_Maddern_2026).

*Statistical analysis*

A three-way analysis of variance (ANOVA) assessed alcohol intake in male and female mice allocated to the alcohol anticipating and binge drinking groups across sessions. A two-tailed unpaired t-test compared total intake (g/kg) between male and female mice during the final 2hr binge drinking test sessions, Fos/mm$^2$, CTβ/mm$^2$ and %CTβ-positive Fos cells between sexes for each BLA efferent projection assessed (mPFC, AcbC, AcbSh, BNST, vHipp), and 0-2secs area under the curve (AUC) between males and females. A two-way ANOVA examined the effect of treatment group (alcohol naïve, alcohol anticipating and binge drinking) and sex on Fos/mm$^2$ expression for each brain region quantified. Inter-regional Pearson's correlations of Fos/mm$^2$ were performed between each brain region quantified for each sex separately for the naïve, alcohol anticipating and binge drinking groups. Repeated measures (RM) two-way ANOVAs were used to assess the effect of treatment (vehicle or CNO) and time (hr) on cumulative alcohol (g/kg) or sucrose (mL) intake during the extended binge drinking test session for each sex (males and females) and viral group (control and DREADD). Two-tailed paired t-tests assessed the effect of treatment (vehicle or CNO) on total alcohol (g/kg) or sucrose (mL) intake in each sex (males and females) and viral group (control and DREADD). One-sample t-tests were used to assess the delta (Δ) change in alcohol intake (g/kg) and sucrose intake (mL) following CNO, relative to saline, administration in control and hM4Di mice. *Post-hoc* Bonferroni corrections for multiple comparisons were performed when statistical significance was achieved. These analyses were performed using GraphPad Prism, with significance being set at $p < 0.05$.

*Post-hoc* fiber photometry analysis was performed using the FiPhoPHA python package (https://pypi.org/project/fiphopha/) (16), with the following settings: 99% confidence interval, 1000 bootstraps, 1000 permutations and a consecutive threshold of 10 datapoints (17).

Data are presented as mean ± standard error mean (SEM).

***Supplementary Figures and Tables:***

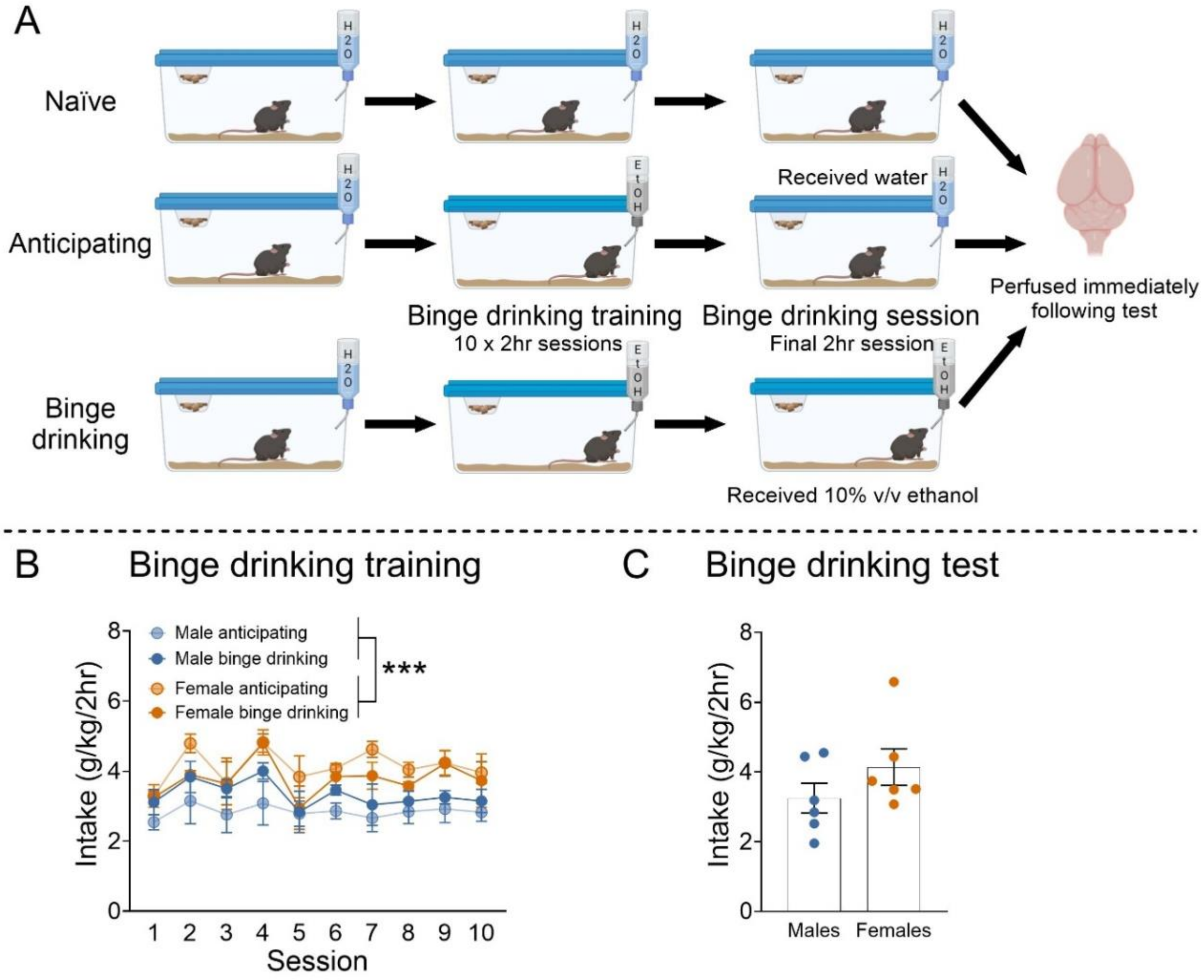


***Supplementary figure 1. Behavioral training data of alcohol anticipating and binge drinking male and female mice. (A)*** *Schematic representation of experimental design.* ***(B)*** *Alcohol intake of male and female mice in the alcohol anticipating and binge drinking groups during binge drinking training sessions, with female mice displaying greater levels of alcohol intake (g/kg) than males (n=6/sex/group).* ***(C)*** *There was no sex difference in total alcohol intake (g/kg) in male and female mice assigned to the binge drinking group during the final binge drinking session. Data presented as mean ± SEM. RM three-way ANOVA, main effect of sex, ***$p<0.001$. ANOVA, analysis of variance, g, gram; hr, hour; kg, kilogram; RM, repeated measures. Please refer to Table S2 for complete statistical analyses.*

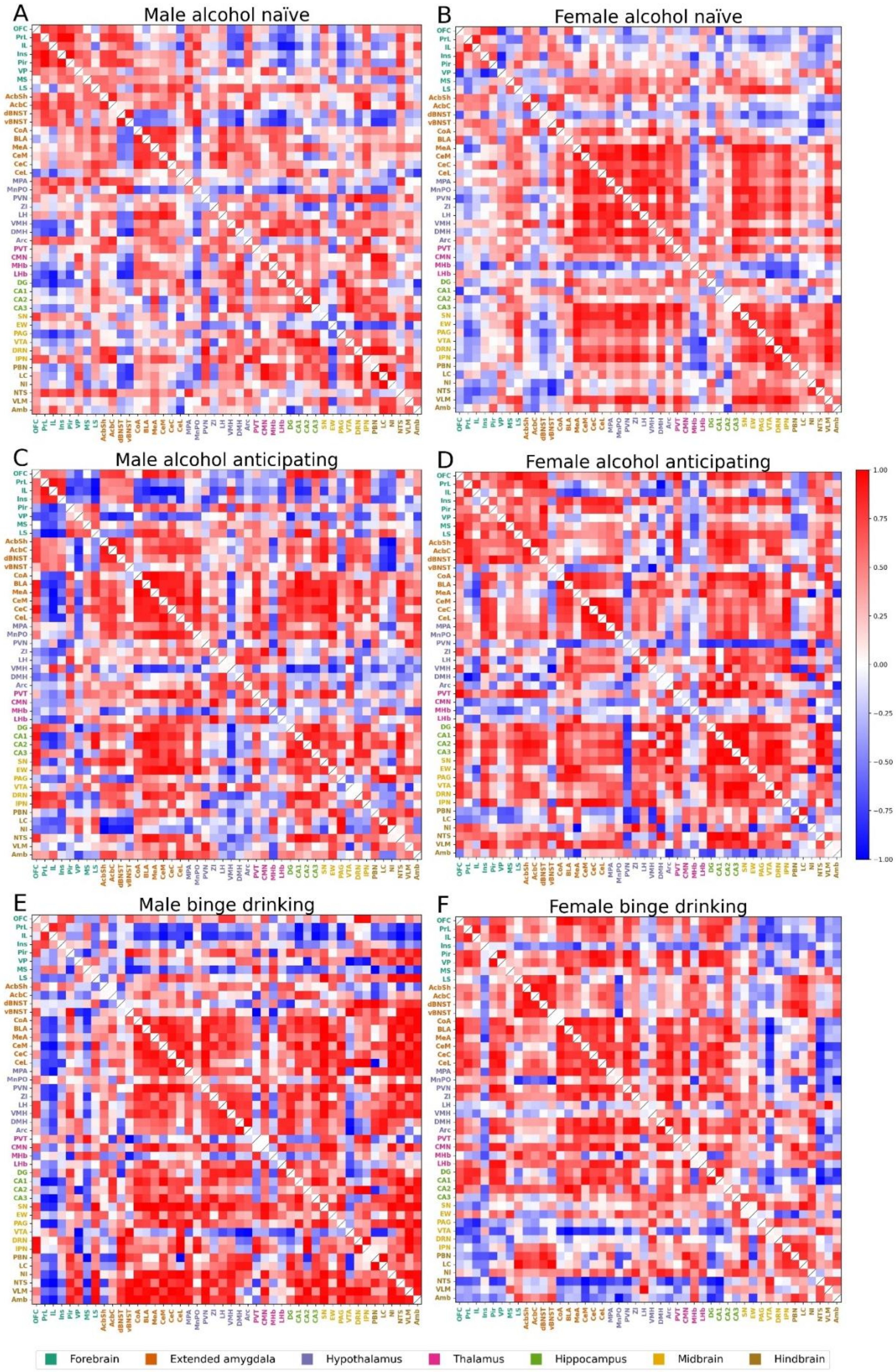
A
Male alcohol naïve
B
Female alcohol naïve
C
Male alcohol anticipating
D
Female alcohol anticipating
E
Male binge drinking
F
Female binge drinking
OFC PrL IL Ins Pir VP MS LS AcbSh AcbC dBNST vBNST CoA BLA MeA CeM CeC CeL MPA MnPO PVN ZI LH VMH DMH Arc PVT CMN MHb LHb DG CA1 CA2 CA3 SN EW PAG VTA DRN IPN PBN LC NI NTS VLM Amb
1.00
0.75
0.50
0.25
0.00
−0.25
−0.50
−0.75
−1.00
Forebrain
Extended amygdala
Hypothalamus
Thalamus
Hippocampus
Midbrain
Hindbrain

***Supplementary figure 2. Inter-regional correlations of c-Fos expression between brain regions for each group.*** *Pearson's correlations were calculated between brain regions to visualize patterns of coordinated activation throughout the brain in male* ***(A)*** *and female* ***(B)*** *alcohol naïve, male* ***(C)*** *and female* ***(D)*** *alcohol anticipating, and male* ***(E)*** *and female* ***(F)*** *binge drinking mice.*

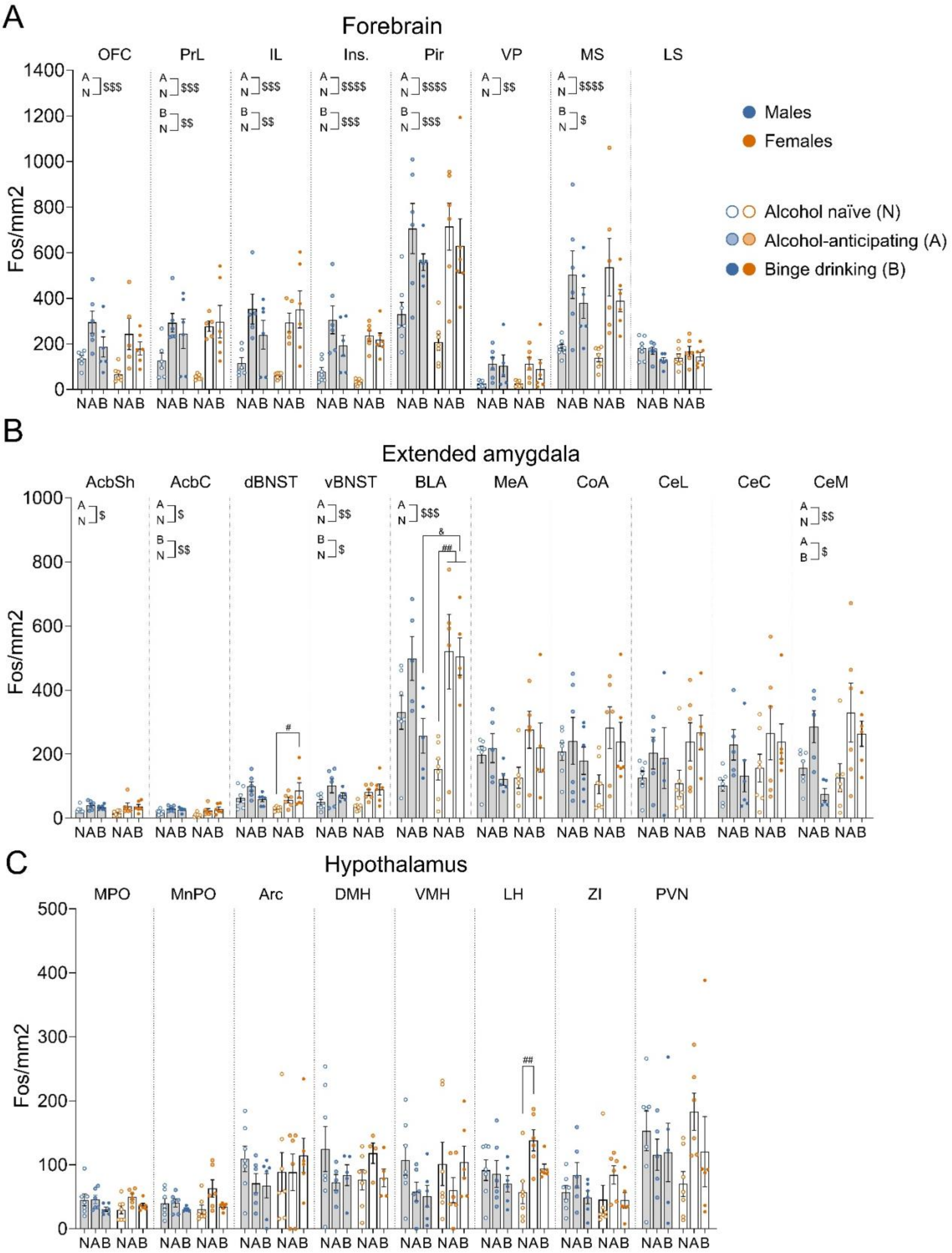

A
Forebrain
OFC
PrL
IL
Ins.
Pir
VP
MS
LS
Fos/mm2
NAB NAB
Males
Females
Alcohol naïve (N)
Alcohol-anticipating (A)
Binge drinking (B)
B
Extended amygdala
AcbSh
AcbC
dBNST
vBNST
BLA
MeA
CoA
CeL
CeC
CeM
C
Hypothalamus
MPO
MnPO
Arc
DMH
VMH
LH
ZI
PVN

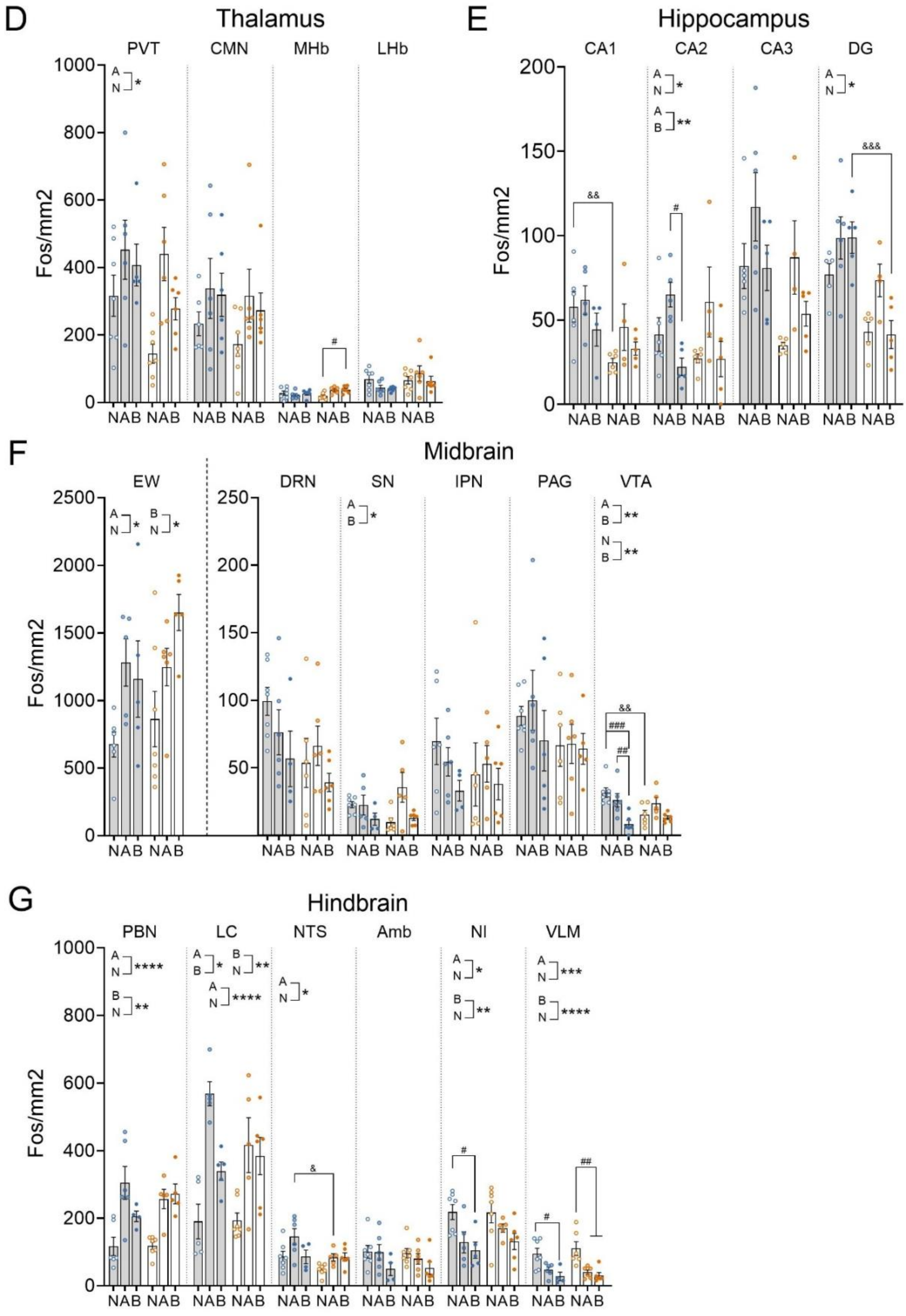

D
Thalamus
PVT
CMN
MHb
LHb
Fos/mm2
NAB NAB
E
Hippocampus
CA1
CA2
CA3
DG
F
Midbrain
EW
DRN
SN
IPN
PAG
VTA
G
Hindbrain
PBN
LC
NTS
Amb
NI
VLM

***Supplementary figure 3. C-Fos expression across the 40 brain regions and subdivisions quantified in naïve, anticipating and binge drinking male and female mice.*** *Quantification of c-Fos expression/mm$^2$ in naïve (N; empty circle), anticipating (A; shaded circle) and binge drinking (B; filled circle) male (blue) and female (orange) mice in brain regions of the* ***(A)*** *forebrain,* ***(B)*** *extended amygdala,* ***(C)*** *hypothalamus,* ***(D)*** *thalamus,* ***(E)*** *hippocampus,* ***(F)*** *midbrain and* ***(H)*** *hindbrain. Data presented as mean ± SEM. Two-way ANOVA assessing sex x treatment group interactions and main effects of sex and treatment group, including Bonferroni post hoc analysis. * = difference between treatment groups regardless of sex ($^{*}p<0.05$, $^{**}p<0.01$, $^{***}p<0.001$, $^{****}p<0.0001$), # = difference between treatment groups within same sex ($^{\#}p<0.05$, $^{\#\#}p<0.01$, $^{\#\#\#}p<0.001$), & = difference between sexes for the same treatment group ($^{\&}p<0.05$, $^{\&\&}p<0.01$, $^{\&\&\&}p<0.001$). n=4-7/sex/group. ANOVA, analysis of variance; A, alcohol anticipating; B, binge drinking; N, naïve. Please refer to Table S1 for brain region abbreviations and Table S5 for complete statistical analyses.*

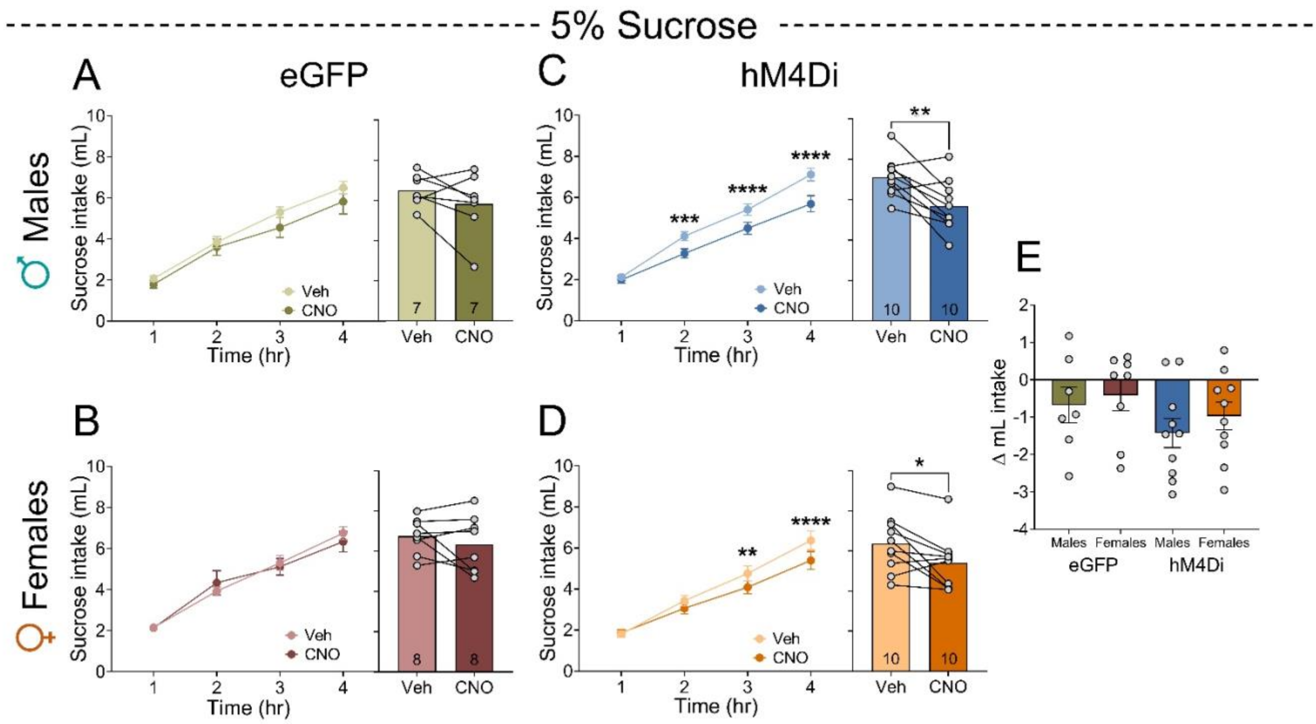


***Supplementary figure 4. Chemogenetic BLA inhibition reduces sucrose intake in male and female mice.*** *CNO administration did not alter cumulative (left) nor total (right) sucrose intake in male* ***(A)*** *or female* ***(B)*** *eGFP control mice (n=7-8/sex). Chemogenetic BLA inhibition reduced both cumulative (left) and total (right) sucrose intake in male* ***(C)*** *and female* ***(D)*** *mice (n=10/sex).* ***(E)*** *No significant difference in Δ sucrose intake following CNO administration was found between eGFP control and hM4Di mice. Data presented as mean ± SEM. *p<0.05, **p<0.01, ***p<0.001, ****p<0.0001. hr, hour; mL, milliliters. Please refer to Table S4 for complete statistical analyses.*

### *Supplementary table 1. Bregma levels quantified across brain regions*

| Brain region | Acronym | Bregma level (mm) |
|---|---|---|
| Orbitofrontal cortex | OFC | +3.20 to +1.98 |
| Prelimbic cortex | PrL | +3.08 to +1.54 |
| Infralimbic cortex | IL | +1.98 to +1.34 |
| Insular cortex | Ins | +2.46 to -1.22 |
| Piriform cortex | Pir | +2.46 to -2.80 |
| Ventral pallidum | VP | +1.94 to -0.34 |
| Medial septum | MS | +1.18 to +0.26 |
| Lateral septum | LS | +1.42 to -0.46 |
| Nucleus accumbens shell/core | AcbSh/AcbC | +1.94 to +0.86 |
| Bed nucleus of the stria terminalis dorsal/ventral | dBNST/vBNST | +0.62 to -0.46 |
| Cortical amygdala | CoA | -0.22 to -3.40 |
| Basolateral amygdala | BLA | -0.58 to -2.06 |
| Medial amygdala | MeA | -0.7 to -2.06 |
| Central amygdala medial division | CeM | -0.58 to -1.58 |
| Central amygdala capsular division | CeC | -0.82 to -1.82 |
| Central amygdala lateral division | CeL | -1.22 to -1.94 |
| Medial preoptic area | MPA | 0.74 to -0.58 |
| Medial preoptic nucleus | MnPO | 0.62 to 0.14 |
| Paraventricular nucleus of the hypothalamus | PVN | -0.58 to -1.34 |
| Zona incerta | ZI | -0.82 to -3.08 |
| Lateral hypothalamus | LH | -0.34 to -2.54 |
| Ventromedial hypothalamus | VMH | -1.06 to -2.06 |
| Dorsomedial hypothalamus | DMH | -1.34 to -2.18 |
| Arcuate nucleus | Arc | -1.46 to -2.06 |
| Paraventricular nucleus of the thalamus | PVT | -0.22 to -1.34 |
| Centromedial nucleus of the thalamus | CMN | -0.46 to -2.06 |
| Medial/lateral habenula | MHb/LHb | -0.94 to -2.18 |
| Dentate gyrus | DG | -0.94 to -4.04 |
| CA1 | CA1 | -1.22 to -3.88 |
| CA2 | CA2 | -1.22 to -3.08 |
| CA3 | CA3 | -0.94 to -3.64 |
| Substantia nigra | SN | -2.46 to -4.04 |
| Edinger-Westphal nucleus | EW | -2.80 to -4.04 |
| Periaqueductal gray | PAG | -2.54 to -5.20 |
| Ventral tegmental area | VTA | -2.92 to -3.88 |
| Dorsal raphe nucleus | DRN | -4.04 to -5.20 |
| Interpeduncular nucleus | IPN | -4.16 to -4.24 |
| Parabrachial nucleus | PBN | -5.02 to -5.68 |
| Locus coeruleus | LC | -5.34 to -5.80 |
| Nucleus incerta | NI | -5.34 to -5.80 |
| Nucleus of the solitary tract | NTS | -6.24 to -7.76 |
| Ventrolateral medulla | VLM | -6.64 to -7.64 |
| Nucleus ambiguous | Amb | -6.64 to -7.92 |

***Supplementary table 2. Fos mapping behavioral data statistical analyses.*** *n=6/group/sex. RM three-way ANOVA. Main effect of session, #p<0.05. Main effect of sex, **p<0.01.*

| | Statistical analysis | Results |
|---|---|---|
| Binge drinking training alcohol intake | RM three-way ANOVA | Session x sex x group interaction: $F_{(4, 80)} = 0.604$, $p = 0.661$<br>Session x sex interaction: $F_{(4, 80)} = 0.431$, $p = 0.786$<br>Session x group interaction: $F_{(4, 80)} = 1.183$, $p = 0.325$<br>Sex x group interaction: $F_{(1, 20)} = 0.103$, $p = 0.752$<br>Main effect of session: $F_{(4, 80)} = 3.561$, $p = 0.010^{\#}$<br>Main effect of sex: $F_{(1, 20)} = 9.422$, $p = 0.006^{**}$<br>Main effect of group: $F_{(1, 20)} = 1.920$, $p = 0.181$ |
| Binge drinking test session alcohol intake | Two-tailed unpaired *t*-test | $t_{(10)} = 1.572$, $p = 0.147$ |

***Supplementary table 3. Global network metrics for each treatment group.***

| *Group* | *Mean Coactivation* | *Anatomical Modularity* | *Community modularity* |
|---|---|---|---|
| Male alcohol naïve | 0.1781 | 0.0379 | 0.3374 |
| Male alcohol anticipating | 0.1672 | 0.0437 | 0.2671 |
| Male binge drinking | 0.2610 | 0.0231 | 0.1986 |
| Female alcohol naïve | 0.2253 | 0.0378 | 0.2238 |
| Female alcohol anticipating | 0.2681 | 0.0090 | 0.1930 |
| Female binge drinking | 0.2080 | 0.0495 | 0.3008 |

***Supplementary table 4. Anatomical partition mean coactivation for each treatment group.*** *ExtAmyg, extended amygdala; Hipp., hippocampus; Hypothal, hypothalamus.*

| *Group* | *Forebrain* | *ExtAmyg* | *Hypothal* | *Thalamus* | *Hipp.* | *Midbrain* | *Hindbrain* |
|---|---|---|---|---|---|---|---|
| Male alcohol naïve | 0.4744 | 0.2553 | 0.1049 | 0.3694 | 0.6593 | 0.0739 | 0.4505 |
| Male alcohol anticipating | -0.0834 | 0.5931 | 0.2503 | 0.1884 | 0.8316 | 0.2521 | 0.3951 |
| Male binge drinking | 0.0257 | 0.3931 | 0.5638 | 0.1607 | 0.5259 | 0.3856 | 0.7179 |
| Female alcohol naïve | 0.0743 | 0.2728 | 0.6639 | 0.3047 | 0.1484 | 0.7794 | 0.3796 |
| Female alcohol anticipating | 0.5248 | 0.3465 | 0.0989 | -0.0498 | 0.9231 | 0.6879 | 0.0207 |
| Female binge drinking | 0.3067 | 0.5408 | 0.4502 | 0.2708 | 0.2666 | 0.2455 | 0.4129 |

***Supplementary table 5. Complete statistical analyses of c-Fos expression for individual brain regions.*** *n=4-7/sex/group. Two-way ANOVA. Treatment group x sex interaction, *p<0.05, **p<0.01. Main effect of treatment group, #p<0.05, ##p<0.01, ###p<0.001, ####p<0.0001. Main effect of sex, &p<0.05, &&p<0.01, &&&&p<0.0001. ANOVA, analysis of variance. Please refer to Supplementary table 1 for brain region acronyms.*

| Region | Treatment x sex interaction | Main effect of treatment | Main effect of sex |
|---|---|---|---|
| OFC | $F_{(2, 30)} = 0.350$, $p = 0.708$ | $F_{(2, 30)} = 9.742$, $p < 0.001$ ### | $F_{(1, 30)} = 1.824$, $p = 0.187$ |
| PrL | $F_{(2, 30)} = 0.964$, $p = 0.393$ | $F_{(2, 30)} = 11.510$, $p <0.001$ ### | $F_{(1, 30)} = 0.104$, $p = 0.750$ |
| IL | $F_{(2, 29)} = 1.760$, $p = 0.190$ | $F_{(2, 29)} = 12.80$, $p < 0.001$ ### | $F_{(1, 29)} = 0.0002$, $p = 0.987$ |
| Ins | $F_{(2, 32)} = 1.042$, $p = 0.364$ | $F_{(2, 32)} = 21.89$, $p < 0.0001$ #### | $F_{(1, 32)} = 1.097$, $p = 0.303$ |
| Pir | $F_{(2, 32)} = 0.786$, $p = 0.464$ | $F_{(2, 32)} = 16.83$, $p < 0.0001$ #### | $F_{(1, 32)} = 0.045$, $p = 0.834$ |
| VP | $F_{(2, 31)} = 0.066$, $p = 0.936$ | $F_{(2, 31)} = 7.927$, $p = 0.002$ ## | $F_{(1, 31)} = 0.342$, $p = 0.563$ |
| MS | $F_{(2, 32)} = 0.158$, $p = 0.855$ | $F_{(2, 32)} = 13.24$, $p < 0.0001$ #### | $F_{(1, 32)} = 5.898$, $p = 0.994$ |
| LS | $F_{(2, 32)} = 1.475$, $p = 0.244$ | $F_{(2, 32)} = 1.740$, $p = 0.192$ | $F_{(1, 32)} = 0.453$, $p = 0.506$ |
| AcbSh | $F_{(2, 32)} = 0.710$, $p = 0.499$ | $F_{(2, 32)} = 5.575$, $p = 0.008$ ## | $F_{(1, 32)} = 0.854$, $p = 0.362$ |
| AcbC | $F_{(2, 32)} = 1.068$, $p = 0.356$ | $F_{(2, 32)} = 6.667$, $p = 0.004$ ## | $F_{(1, 32)} = 1.777$, $p = 0.192$ |
| dBNST | $F_{(2, 32)} = 4.187$, $p = 0.024$* | $F_{(2, 32)} = 3.599$, $p = 0.039$ # | $F_{(1, 32)} = 2.258$, $p = 0.143$ |
| vBNST | $F_{(2, 31)} = 1.246$, $p = 0.302$ | $F_{(2, 31)} = 8.078$, $p = 0.002$ ## | $F_{(1, 31)} = 0.230$, $p = 0.635$ |
| CoA | $F_{(2, 32)} = 1.744$, $p = 0.191$ | $F_{(2, 32)} = 2.238$, $p = 0.123$ | $F_{(1, 32)} = 0.004$, $p = 0.950$ |
| BLA | $F_{(2, 28)} = 5.705$, $p = 0.008$** | $F_{(2, 28)} = 9.099$, $p < 0.001$ ### | $F_{(1, 28)} = 0.331$, $p = 0.570$ |
| MeA | $F_{(2, 27)} = 2.124$, $p = 0.139$ | $F_{(2, 27)} = 2.131$, $p = 0.138$ | $F_{(1, 27)} = 0.560$, $p = 0.447$ |
| CeM | $F_{(2, 30)} = 2.450$, $p = 0.103$ | $F_{(2, 30)} = 6.034$, $p = 0.006$ ## | $F_{(1, 30)} = 2.640$, $p = 0.115$ |
| CeC | $F_{(2, 31)} = 0.233$, $p = 0.794$ | $F_{(2, 31)} = 2.609$, $p = 0.090$ | $F_{(1, 31)} = 2.396$, $p = 0.132$ |
| CeL | $F_{(2, 28)} = 0.463$, $p = 0.634$ | $F_{(2, 28)} = 3.187$, $p = 0.057$ | $F_{(1, 28)} = 0.591$, $p = 0.448$ |
| MPA | $F_{(2, 32)} = 1.877$, $p = 0.170$ | $F_{(2, 32)} = 2.724$, $p = 0.081$ | $F_{(1, 32)} = 0.108$, $p = 0.745$ |
| MnPO | $F_{(2, 32)} = 2.297$, $p = 0.117$ | $F_{(2, 32)} = 3.619$, $p = 0.038$ # | $F_{(1, 32)} = 0.864$, $p = 0.360$ |
| PVN | $F_{(2, 30)} = 2.337$, $p = 0.114$ | $F_{(2, 30)} = 0.609$, $p = 0.551$ | $F_{(1, 30)} = 0.025$, $p = 0.875$ |
| ZI | $F_{(2, 32)} = 0.076$, $p = 0.927$ | $F_{(2, 32)} = 3.075$, $p = 0.060$ | $F_{(1, 32)} = 0.170$, $p = 0.683$ |
| LH | $F_{(2, 32)} = 4.006$, $p = 0.028$* | $F_{(2, 32)} = 3.067$, $p = 0.061$ | $F_{(1, 32)} = 1.123$, $p = 0.297$ |
| VMH | $F_{(2, 32)} = 0.886$, $p = 0.422$ | $F_{(2, 32)} = 1.840$, $p = 0.175$ | $F_{(1, 32)} = 0.716$, $p = 0.404$ |
| DMH | $F_{(2, 27)} = 2.083$, $p = 0.144$ | $F_{(2, 27)} = 0.377$, $p = 0.690$ | $F_{(1, 27)} = 0.014$, $p = 0.908$ |
| Arc | $F_{(2, 32)} = 1.008$, $p = 0.376$ | $F_{(2, 32)} = 0.328$, $p = 0.723$ | $F_{(1, 32)} = 0.560$, $p = 0.460$ |
| PVT | $F_{(2, 31)} = 0.921$, $p = 0.409$ | $F_{(2, 31)} = 6.522$, $p = 0.004$ ## | $F_{(1, 31)} = 4.310$, $p = 0.046$ & |
| CMN | $F_{(2, 31)} = 0.054$, $p = 0.947$ | $F_{(2, 31)} = 2.289$, $p = 0.118$ | $F_{(1, 31)} = 0.741$, $p = 0.396$ |
| MHb | $F_{(2, 32)} = 3.916$, $p = 0.030$* | $F_{(2, 32)} = 1.962$, $p = 0.157$ | $F_{(1, 32)} = 1.990$, $p = 0.168$ |
| LHb | $F_{(2, 32)} = 1.550$, $p = 0.228$ | $F_{(2, 32)} = 0.826$, $p = 0.447$ | $F_{(1, 32)} = 3.687$, $p = 0.064$ |
| DG | $F_{(2, 24)} = 1.573$, $p = 0.228$ | $F_{(2, 24)} = 3.883$, $p = 0.035$ # | $F_{(1, 24)} = 25.49$, $p < 0.0001$ &&&& |
| CA1 | $F_{(2, 24)} = 1.049$, $p = 0.366$ | $F_{(2, 24)} = 1.815$, $p = 0.185$ | $F_{(1, 24)} = 8.999$, $p = 0.006$ && |
| CA2 | $F_{(2, 25)} = 0.500$, $p = 0.613$ | $F_{(2, 25)} = 7.639$, $p = 0.003$ ## | $F_{(1, 25)} = 0.341$, $p = 0.565$ |
| CA3 | $F_{(2, 25)} = 0.273$, $p = 0.763$ | $F_{(2, 25)} = 4.733$, $p = 0.018$ | $F_{(1, 25)} = 8.204$, $p = 0.008$ && |
| SN | $F_{(2, 27)} = 2.988$, $p = 0.067$ | $F_{(2, 327)} = 4.678$, $p = 0.018$ # | $F_{(1, 27)} = 0.003$, $p = 0.954$ |
| EW | $F_{(2, 28)} = 0.981$, $p = 0.387$ | $F_{(2, 28)} = 7.112$, $p = 0.003$ ## | $F_{(1, 28)} = 2.094$, $p = 0.159$ |

| | | | |
|---|---|---|---|
| PAG | $F_{(2, 30)} = 0.313$, $p = 0.734$ | $F_{(2, 30)} = 0.511$, $p = 0.605$ | $F_{(1, 30)} = 2.261$, $p = 0.143$ |
| VTA | $F_{(2, 30)} = 4.873$, $p = 0.015$* | $F_{(2, 30)} = 9.474$, $p < 0.001^{###}$ | $F_{(1, 30)} = 2.615$, $p = 0.116$ |
| DRN | $F_{(2, 29)} = 0.880$, $p = 0.426$ | $F_{(2, 29)} = 1.977$, $p = 0.157$ | $F_{(1, 29)} = 4.105$, $p = 0.052$ |
| IPN | $F_{(2, 27)} = 0.485$, $p = 0.621$ | $F_{(2, 27)} = 1.022$, $p = 0.373$ | $F_{(1, 27)} = 0.289$, $p = 0.595$ |
| PBN | $F_{(2, 27)} = 1.610$, $p = 0.218$ | $F_{(2, 27)} = 17.21$, $p < 0.0001^{####}$ | $F_{(1, 27)} = 0.078$, $p = 0.782$ |
| LC | $F_{(2, 27)} = 2.241$, $p = 0.126$ | $F_{(2, 27)} = 19.79$, $p < 0.0001^{####}$ | $F_{(1, 27)} = 0.778$, $p = 0.386$ |
| NI | $F_{(2, 29)} = 0.381$, $p = 0.687$ | $F_{(2, 29)} = 8.393$, $p = 0.001^{##}$ | $F_{(1, 29)} = 1.093$, $p = 0.304$ |
| NTS | $F_{(2, 29)} = 1.775$, $p = 0.187$ | $F_{(2, 29)} = 4.965$, $p = 0.014^{#}$ | $F_{(1, 29)} = 6.894$, $p = 0.014^{\&}$ |
| VLM | $F_{(2, 29)} = 0.375$, $p = 0.691$ | $F_{(2, 29)} = 16.73$, $p < 0.0001^{####}$ | $F_{(1, 29)} = 0.091$, $p = 0.765$ |
| Amb | $F_{(2, 30)} = 0.179$, $p = 0.837$ | $F_{(2, 30)} = 3.360$, $p = 0.048^{#}$ | $F_{(1, 30)} = 0.194$, $p = 0.663$ |

***Supplementary table 6. In vivo fiber photometry statistical analyses.*** *n*=6/sex. Two-tailed *unpaired t-test, *p<0.05.*

| Data analyzed (two-tailed unpaired *t*-test) | Results |
|---|---|
| Alcohol AUC 0-2secs from onset of drinking | $t_{(10)} = 2.243$, $p = 0.049$* |
| Sucrose AUC 0-2ses from onset of drinking | $t_{(10)} = 0.733$, $p = 0.480$ |

***Supplementary table 7. Chemogenetic BLA inhibition statistical analyses.*** *n=35 (males 10 DREADD, 7 eGFP control; females 10 DREADD, 8 eGFP control). RM two-way ANOVA. Treatment x time interaction, $^{\&}p<0.05$, $^{\&\&}p<0.01$, $^{\&\&\&}p<0.001$. Main effect of treatment, *p<0.05, **p<0.01. Main effect of time, $^{####}p<0.0001$. Main effect of viral group, *p<0.05. Two-tailed paired t-test, *p<0.05, **p<0.01. One-sample t-test, *p<0.05, **p<0.01.*

| Data analyzed | Statistical analysis | Results |
|---|---|---|
| Male eGFP cumulative ethanol intake | RM two-way ANOVA | Treatment X time interaction: $F_{(3, 18)} = 3.074$, $p = 0.054$<br>Main effect of treatment: $F_{(1, 6)} = 0.694$, $p = 0.437$<br>Main effect of time: $F_{(3, 18)} = 122.4$, $p < 0.0001^{####}$ |
| Male eGFP total ethanol intake | Two-tailed paired *t*-test | $t_{(6)} = 0.255$, $p = 0.807$ |
| Female eGFP cumulative ethanol intake | RM two-way ANOVA | Treatment X time interaction: $F_{(3, 21)} = 0.330$, $p = 0.804$<br>Main effect of treatment: $F_{(1, 7)} = 0.218$, $p = 0.655$<br>Main effect of time: $F_{(3, 21)} = 160.5$, $p < 0.0001^{####}$ |
| Female eGFP total ethanol intake | Two-tailed paired *t*-test | $t_{(7)} = 0.047$, $p = 0.974$ |

| | | |
|---|---|---|
| Male hM4Di cumulative ethanol intake | RM two-way ANOVA | Treatment X time interaction: $F_{(3, 27)} = 1.909$, $p = 0.152$<br>Main effect of treatment: $F_{(1, 9)} = 10.16$, $p = 0.011$*<br>Main effect of time: $F_{(3, 27)} = 163.8$, $p < 0.0001^{\#\#\#\#}$ |
| Male hM4Di total ethanol intake | Two-tailed paired *t*-test | $t_{(9)} = 2.186$, $p = 0.057$ |
| Female hM4Di cumulative ethanol intake | RM two-way ANOVA | Treatment X time interaction: $F_{(3, 27)} = 3.495$, $p = 0.029^{\&}$<br>Main effect of treatment: $F_{(1, 9)} = 9.836$, $p = 0.012$*<br>Main effect of time: $F_{(3, 18)} = 74.45$, $p < 0.0001^{\#\#\#\#}$ |
| Female hM4Di total ethanol intake | Two-tailed paired *t*-test | $t_{(9)} = 3.883$, $p = 0.004^{**}$ |
| Delta change in alcohol intake following CNO | Two-way ANOVA | Viral group X sex interaction: $F_{(1, 31)} = 0.045$, $p = 0.833$<br>Main effect of viral group: $F_{(1, 31)} = 7.482$, $p = 0.010$*<br>Main effect of sex: $F_{(1, 31)} = 0.137$, $p = 0.714$ |
| Male eGFP cumulative sucrose intake | RM two-way ANOVA | Treatment X time interaction: $F_{(3, 18)} = 1.053$, $p = 0.379$<br>Main effect of treatment: $F_{(1, 6)} = 1.895$, $p = 0.218$<br>Main effect of time: $F_{(3, 18)} = 168.1$, $p < 0.0001^{\#\#\#\#}$ |
| Male eGFP total sucrose intake | Two-tailed paired *t*-test | $t_{(6)} = 1.408$, $p = 0.209$ |
| Female eGFP cumulative sucrose intake | RM two-way ANOVA | Treatment X time interaction: $F_{(3, 21)} = 1.229$, $p = 0.322$<br>Main effect of treatment: $F_{(1, 7)} = 0.023$, $p = 0.884$<br>Main effect of time: $F_{(3, 21)} = 113.1$, $p < 0.0001^{\#\#\#\#}$ |
| Female eGFP total sucrose intake | Two-tailed paired *t*-test | $t_{(7)} = 1.001$, $p = 0.350$ |
| Male hM4Di cumulative sucrose intake | RM two-way ANOVA | Treatment X time interaction: $F_{(3, 27)} = 9.145$, $p < 0.001^{\&\&\&}$<br>Main effect of treatment: $F_{(1, 9)} = 10.6$, $p = 0.010$**<br>Main effect of time: $F_{(3, 27)} = 291.9$, $p < 0.0001^{\#\#\#\#}$ |
| Male hM4Di total sucrose intake | Two-tailed paired *t*-test | $t_{(9)} = 3.640$, $p = 0.005$** |
| Female hM4Di cumulative sucrose intake | RM two-way ANOVA | Treatment X time interaction: $F_{(3, 27)} = 6.286$, $p = 0.002^{\&\&}$<br>Main effect of treatment: $F_{(1, 9)} = 6.644$, $p = 0.030$*<br>Main effect of time: $F_{(3, 27)} = 151.1$, $p < 0.0001^{\#\#\#\#}$ |
| Female hM4Di total sucrose intake | Two-tailed paired *t*-test | $t_{(9)} = 2.617$, $p = 0.028$* |
| Delta change in sucrose intake following CNO | Two-way ANOVA | Viral group X sex interaction: $F_{(1, 31)} = 0.052$, $p = 0.820$<br>Main effect of viral group: $F_{(1, 31)} = 2.479$, $p = 0.126$<br>Main effect of sex: $F_{(1, 31)} = 0.741$, $p = 0.396$ |

***Supplementary table 8. Activation of BLA efferent projections following binge drinking statistical analyses.*** *n=24/sex (mPFC, n=5/sex; AcbC, n=5/sex; AcbSh, n=8/sex; BNST, n=6 males, 7 females; vHipp, n=5 males, 8 females). Two-tailed unpaired t-test, *p<0.05, **p<0.01.*

| Data analysed (two-tailed unpaired *t*-test) | Results |
|---|---|
| Fos/mm$^2$ | $t_{(46)} = 3.021$, $p = 0.004$** |
| BLA→mPFC | Fos: $t_{(8)} = 2.384$, $p = 0.044$*<br>CTβ: $t_{(8)} = 3.350$, $p = 0.010$*<br>%CTβ-positive Fos cells: $t_{(8)} = 3.400$, $p=0.009$** |
| BLA→AcbC | Fos: $t_{(8)} = 2.587$, $p = 0.032$*<br>CTβ: $t_{(8)} = 2.664$, $p = 0.029$*<br>%CTβ-positive Fos cells: $t_{(8)} = 4.191$, $p = 0.003$** |
| BLA→AcbSh | Fos: $t_{(14)} = 0.683$, $p = 0.506$<br>CTβ: $t_{(14)} = 0.052$, $p = 0.959$<br>%CTβ-positive Fos cells: $t_{(14)} = 0.852$, $p = 0.409$ |
| BLA→BNST | Fos: $t_{(11)} = 1.742$, $p = 0.109$<br>CTβ: $t_{(11)} = 0.506$, $p = 0.623$<br>%CTβ-positive Fos cells: $t_{(11)} = 0.109$, $p = 0.915$ |
| BLA→vHipp | Fos: $t_{(11)} = 2.366$, $p = 0.037$*<br>CTβ: $t_{(11)} = 0.383$, $p = 0.709$<br>%CTβ-positive Fos cells: $t_{(11)} = 0.508$, $p = 0.621$ |

***Supplementary table 9. BLA→AcbC pathway-specific inhibition statistical analyses.*** *n=16 (7 males, 9 females). RM two-way ANOVA. Treatment x time interaction, $^{\&\&}p<0.01$. Main effect of time, $^{\#\#\#\#}p<0.0001$. Two-tailed paired t-test, *p<0.05. One-sample t-test, *p<0.05.*

| Data analyzed | Statistical analysis | Results |
|---|---|---|
| Male cumulative ethanol intake | RM two-way ANOVA | Treatment X time interaction: $F_{(3, 18)} = 1.133$, $p = 0.362$<br>Main effect of treatment: $F_{(1, 6)} = 0.651$, $p = 0.451$<br>Main effect of time: $F_{(3, 18)} = 80.42$, $p < 0.0001^{\#\#\#\#}$ |
| Male total ethanol intake | Two-tailed paired *t*-test | $t_{(6)} = 1.338$, $p = 0.230$ |
| Female cumulative ethanol intake | RM two-way ANOVA | Treatment X time interaction: $F_{(3, 24)} = 6.148$, $p = 0.003^{\&\&}$<br>Main effect of treatment: $F_{(1, 8)} = 4.859$, $p = 0.059$<br>Main effect of time: $F_{(3, 24)} = 158.7$, $p < 0.0001^{\#\#\#\#}$ |
| Female total ethanol intake | Two-tailed paired *t*-test | $t_{(8)} = 2.345$, $p = 0.047$* |
| Male cumulative sucrose intake | RM two-way ANOVA | Treatment X time interaction: $F_{(3, 18)} = 0.440$, $p = 0.727$<br>Main effect of treatment: $F_{(1, 6)} = 4.336$, $p = 0.083$<br>Main effect of time: $F_{(3, 18)} = 240$, $p < 0.0001^{\#\#\#\#}$ |
| Male total sucrose intake | Two-tailed paired *t*-test | $t_{(6)} = 1.654$, $p = 0.149$ |
| Female cumulative sucrose intake | RM two-way ANOVA | Treatment X time interaction: $F_{(3, 24)} = 1.714$, $p = 0.191$<br>Main effect of treatment: $F_{(1, 8)} = 0.7586$, $p = 0.409$<br>Main effect of time: $F_{(3, 24)} = 277.0$, $p < 0.0001^{\#\#\#\#}$ |

| | | |
|---|---|---|
| Female total sucrose intake | Two-tailed paired *t*-test | $t_{(8)} = 1.145$, $p = 0.285$ |
| Delta change in ethanol intake following CNO | Two-tailed unpaired *t*-test | $t_{(14)} = 2.524$, $p = 0.023$* |
| Delta change in sucrose intake following CNO | Two-tailed unpaired *t*-test | $t_{(14)} = 0.296$, $p = 0.772$ |
